# 14-3-3ζ constrains insulin secretion by regulating mitochondrial function in pancreatic β-cells

**DOI:** 10.1101/2021.10.17.464702

**Authors:** Y Mugabo, C Zhao, JJ Tan, A Ghosh, SA Campbell, E Fadzeyeva, F Paré, SS Pan, M Galipeau, J Ast, J Broichhagen, DJ Hodson, EE Mulvihill, S Petropoulos, GE Lim

## Abstract

While critical for neurotransmitter synthesis in the brain, members of the 14-3-3 protein family are often assumed to have redundant, over-lapping roles due to their high sequence homology and ubiquitous expression. Despite this assumption, various mammalian 14-3-3 isoforms have now been implicated in regulating cellular and organismal metabolism; however, these functions were primarily observed in cell lines or from systemic knockout mouse models. To date, we have begun to define the contributions of 14-3-3ζ in adipocytes, but whether 14-3-3ζ has additional metabolic roles in other cell types, such as the pancreatic β-cell, is unclear. We previously documented a pro-survival role of 14-3-3ζ in MIN6 insulinoma cells, as depletion of 14-3-3ζ induced cell death, but paradoxically, whole-body deletion of 14-3-3ζ in mice resulted in significantly enlarged β-cell area with no effects on insulin secretion. To better understand the role of 14-3-3ζ in β-cells, we generated β-cell-specific 14-3-3ζ knockout (β14-3-3ζKO) mice, and while no differences in β-cell mass were observed, β14-3-3ζKO mice displayed potentiated insulin secretion due to enhanced mitochondrial function and ATP synthesis. Deletion of 14-3-3ζ led to profound changes to the β-cell transcriptome, where pathways associated with mitochondrial respiration and oxidative phosphorylation were upregulated. Acute treatment of mouse islets and human islets with pan-14-3-3 inhibitors recapitulated the potentiation in glucose-stimulated insulin secretion (GSIS) and mitochondrial function, suggesting that 14-3-3ζ is a critical isoform in β-cells that regulates GSIS. In dysfunctional *db/db* islets and islets from type 2 diabetic donors, expression of *Ywhaz*/*YWHAZ*, the gene encoding 14-3-3ζ, was inversely associated with insulin secretory capacity, and pan-14-3-3 protein inhibition was capable of enhancing GSIS and mitochondrial function. Taken together, this study demonstrates important regulatory functions of 14-3-3ζ and its related isoforms in insulin secretion and mitochondrial function in β-cells. A deeper understanding of how 14-3-3ζ influences β-cell function will further advance our knowledge of how insulin secretion from β-cells is regulated.

## Introduction

The downstream signaling events underlying glucose-stimulated insulin secretion (GSIS) are well-conserved in rodent and human β-cells. Following the uptake of glucose into pancreatic β-cells by low-affinity transporters, it is converted into pyruvate before entering the tricarboxylic acid (TCA) cycle and oxidative phosphorylation cascade to yield ATP (Hou et al., 2009). Increases in the ATP/ADP ratio then lead to closure of K-ATP channels to depolarize the β-cell and the subsequent opening of voltage-dependent Ca^2+^ channels. Along with amplifying signals, the rise in intracellular Ca^2+^drive insulin granule exocytosis. Insulin secretion follows two phases: a rapid, maximal release period of insulin release, followed by a lower magnitude, sustained second-phase of insulin secretion (Henquin, 2021). Aside from closing K-ATP channels, ATP is also required for kinesin-mediated translocation of latent insulin granules to the plasma membrane for the second-phase of insulin secretion (Rorsman and Renstrom, 2003). These events require precise spatial and temporal coordination to ensure proper insulin release to maintain normoglycemia (Almaca et al., 2015).

Scaffold proteins are essential regulators of signaling events due to their ability to facilitate the translocation of downstream effectors and influence the activities of receptors, kinases, and enzymes (Diallo et al., 2019; Mugabo and Lim, 2018). In β-cells, various scaffolds, such as β-arrestin-2, AKAP150, and NCK1, have been reported to be necessary for insulin secretion (Hinke et al., 2012; Yamani et al., 2015; Zhu et al., 2017). For example, β-cell-specific deletion or siRNA-mediated knockdown of β-arrestin-2 in mouse β-cells or EndoC-βH1 cells, respectively, leads to defective GSIS (Zhu et al., 2017). Additionally, islets from systemic AKAP150 knockout mice display impaired insulin secretion due to alterations in Ca^2+^-evoked currents (Hinke et al., 2012).

14-3-3 proteins are a ubiquitously expressed family of scaffolds that were initially discovered in brain extracts, and they comprise approximately 1% of soluble proteins in the brain (Diallo et al., 2019; Mugabo and Lim, 2018). Their ability to recognize serine- or threonine phosphorylated proteins results in a large interactome that is shared among all seven mammalian isoforms (Dubois et al., 2009; Jin et al., 2004; Mugabo et al., 2018). Despite exhibiting high sequence homology, isoform-specific roles of 14-3-3 proteins in various cell types, including insulinoma cells and adipocytes, have been identified (Lim et al., 2015; Lim et al., 2013). We previously identified 14-3-3ζ as a critical regulator of adipogenesis, glucose homeostasis, and pancreatic β-cell survival (Lim et al., 2015; Lim and Johnson, 2016; Lim et al., 2013; Lim et al., 2016). Systemic deletion of 14-3-3ζ in mice was associated with significant reductions in adipogenesis, and siRNA-mediated depletion of only 14-3-3ζ in 3T3-L1 pre-adipocytes abrogated adipocyte differentiation (Lim et al., 2015). With respect to glucose homeostasis, whole-body 14-3-3ζ knockout mice displayed enhanced oral glucose tolerance due to significantly elevated fasting levels of the incretin hormone, GLP-1 (Lim et al., 2016).

One of the earliest indications of 14-3-3 proteins being involved in hormone secretion was identified by Morgan and Burgoyne, who found that 14-3-3 proteins participate in calcium-dependent release of catecholamines from bovine adrenal chromaffin cells (Morgan and Burgoyne, 1992). The contribution of 14-3-3 proteins in exocytosis is also conserved across organisms, as the Drosophila homolog of 14-3-3ζ, Leonardo, was discovered to regulate the dynamics of synaptic vesicles and synaptic transmission rates (Broadie et al., 1997). We first explored the possibility of 14-3-3 proteins to influence insulin secretion using MIN6 insulinoma cells, and plasmid-based over-expression of difopein, a 14-3-3 protein inhibitor, in dispersed mouse islets or MIN6 cells attenuated insulin secretion, but this was also associated with increased cell death, an event known to be associated with impaired insulin secretion (Akerfeldt et al., 2008; Bernal-Mizrachi et al., 2004; Carrington et al., 2009; Lim et al., 2013). Depletion of individual isoforms by siRNA in MIN6 cells had no impact on insulin secretion, whereas over-expression of HA-14-3-3ζ attenuated GSIS (Lim et al., 2016). Moreover, siRNA-mediated knockdown and over-expression of 14-3-3ζ were found to complimentarily induce MIN6 cell apoptosis or promote survival, respectively (Lim et al., 2013). Interestingly systemic 14-3-3ζ knockout mice had increased β-cell area, which suggested proliferative actions of 14-3-3ζ in the β-cell (Lim et al., 2016). With these conflicting results from MIN6 insulinoma cells and whole-body 14-3-3ζKO mice, it is unclear whether 14-3-3ζ has cell-autonomous roles in β-cells to influence insulin secretion, cell survival, or proliferation. Thus, in-depth studies are required to truly understand the β-cell specific roles of 14-3-3ζ.

Using a combination of approaches and models, we identify 14-3-3ζ as a critical regulator of insulin secretion in primary β-cells. Deletion of 14-3-3ζ specifically in β-cells enhanced GSIS, demonstrating a physiological role of 14-3-3ζ to constrain insulin secretion. Single-cell RNA-seq revealed significant upregulation of genes and pathways associated with mitochondrial respiration and oxidative phosphorylation in 14-3-3-deficient β-cells, which were reflected by changes in mitochondrial mass and activity. Acute inhibition of all 14-3-3 proteins in primary mouse and human islets recapitulated the potentiation of GSIS and mitochondrial function that occurred following 14-3-3ζ deletion in β-cells, suggesting that 14-3-3ζ could be a key isoform in human β-cells. Expression levels of *Ywhaz/YWHAZ,* the gene encoding 14-3-3ζ, were found to be inversely associated with insulin secretory capacity and significantly elevated in dysfunctional *db/db* islets and human islets from type 2 diabetic donors, and inhibition of 14-3-3 proteins was able to alleviate β-cell dysfunction. Taken together, this study unequivocally demonstrates the ability of 14-3-3ζ to restrain insulin release from pancreatic β-cells by influencing mitochondrial mass and function. Moreover, it further deepens our knowledge of the regulatory factors present in a β-cell that control insulin secretion.

## Results

### Inhibition of 14-3-3 proteins is sufficient to increase insulin secretion

We previously reported that over-expression of the 14-3-3 protein peptide inhibitor, difopein, in MIN6 cells and dispersed mouse islets impaired GSIS, in addition to inducing cell death (Lim et al., 2013; Masters and Fu, 2001). Given the importance of cell-cell contact in propagating signals underlying insulin secretion (Johnston et al., 2016; Konstantinova et al., 2007) and differences in insulin secretion between cell lines and primary β-cells (Schulze et al., 2016), intact mouse and human islets were treated with cell-permeable 14-3-3 inhibitors, 14-3-3i and BV02, both of which potentiated GSIS (Fig. 1A, I). Opening of K-ATP channels with diazoxide was sufficient to prevent the potentiation of GSIS due to 14-3-3 inhibitors, demonstrating that the effects of 14-3-3 protein inhibition on GSIS were upstream to that of K-ATP channel closure in mouse and human islets (Fig. 1A, I). To address whether inhibition of 14-3-3 proteins altered the responsiveness of β-cells to glucose, normal mouse islets were exposed to different concentrations of glucose in the absence or presence of 14-3-3i or BV02. Inhibition of 14-3-3 proteins lowered the glucose threshold to stimulate GSIS such that 10 mM glucose was sufficient to stimulate insulin secretion (Suppl Fig. 1A). Potentiated GSIS was observed following 14-3-3 protein inhibition at each test concentration of glucose, but no concentration of glucose was able to exceed the maximal amount of secreted insulin in response to 25 mM glucose (Suppl Fig. 1A).

**Figure 1.**
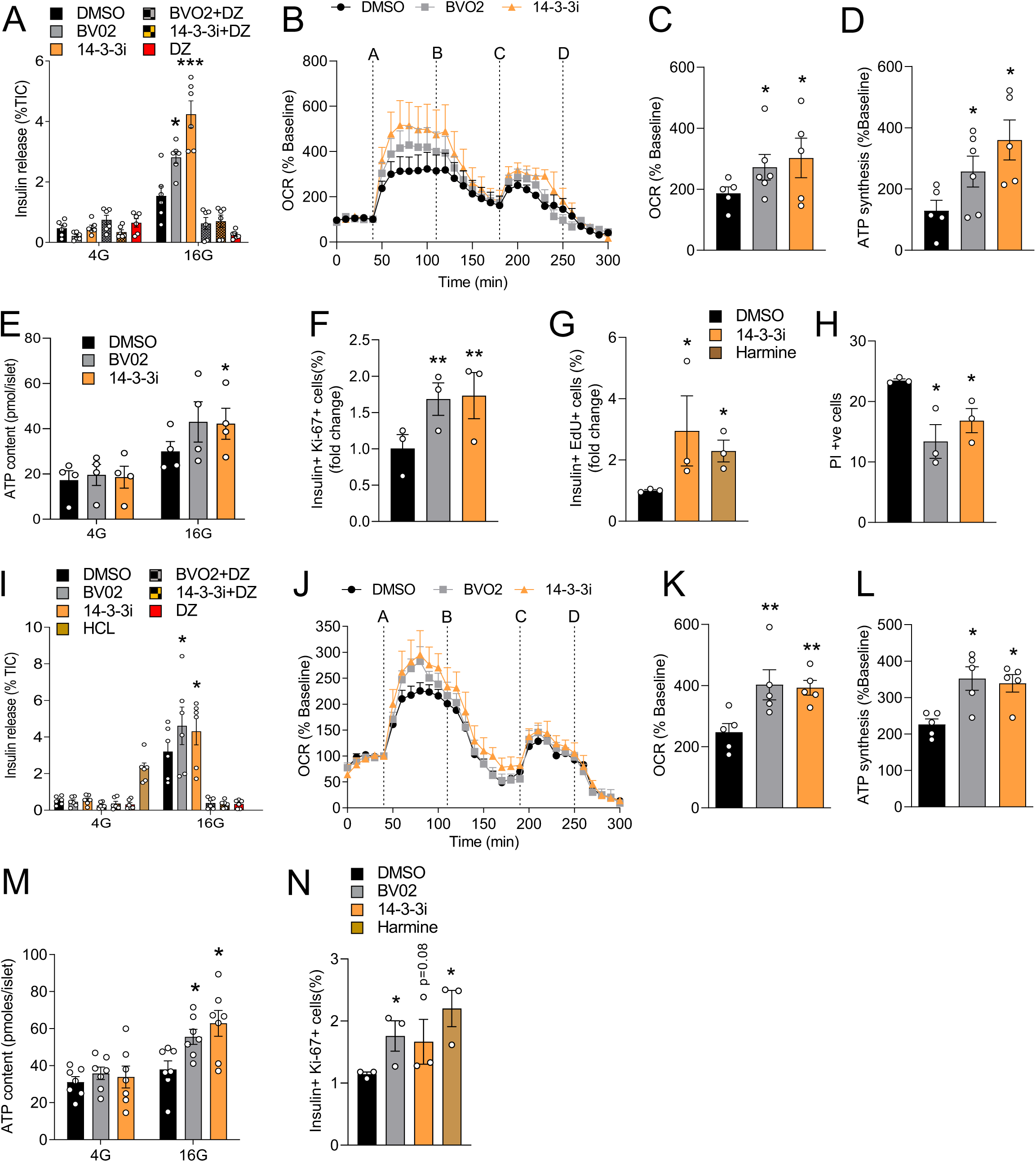
Pharmacological inhibition of 14-3-3 proteins in pancreatic islets enhances insulin secretion, mitochondrial function, and proliferation. Mouse **(A-H)** and human islets **(I-N)** were incubated with two pan-14-3-3 inhibitors (14-3-3i and BVO2; 10 μM each) plus or minus diazoxide (DZ, 200μM) for 1hr at 4 mM glucose prior to treatment with 4 (4G) or 16 (16G) mM glucose for 1hr. Insulin secretion (A and I) was measured by RIA and normalized to total insulin content (n=5-6 mice or donors per group; *: p<0.05, ***: p<0.001 when compared to DMSO 16G). Oxygen consumption (B, C, J, K) and ATP synthesis rates (D,L) were measured in mouse (B-D; n=5-6 mice per group; *: p<0.05 when compared to DMSO) and human (J-L; n=5-6 donors; *:p<0.05 when compared to DMSO, **: p<0.01 when compared to DMSO) islets treated with 14-3-3 inhibitors (In panels B and J, oxygen consumption rate (OCR) profiles in response to *(A)* Glucose (16 mM) stimulation*, (B)* Oligomycin (5μM) -mediation inhibition of ATP synthase, *(C)* Uncoupling of respiration due to FCCP (1μM), and (D) inhibition of Complex I and III by *(D)* rotenone (5μM) and antimycin (5μM) are shown). Glucose-stimulated ATP synthesis rates (D,L) were calculated by measuring the decrease in OCR upon injection of oligomycin. (E,M) Biochemical measurements of ATP content in isolated mouse (E; n=4 per group; *: p<0.05) and human (M; n= 7 donors; *: p<0.05) islets treated with 14-3-3 inhibitors and quantified at different glucose concentrations. (F,N) In dispersed mouse (F; n=3 per group; *: p<0.05) and human (N; n=3 donors; *p<0.05) islet preparations, β-cell proliferation, as measured by immunofluorescent staining for Insulin- and Ki67-positive β-cells, was measured after 72 hr treatment with DMSO, 14-3-3 inhibitors (10 μM each), or Harmine (10μM). (G) Mouse β-cell proliferation was quantified by flow cytometry-mediated detection of Insulin- and EdU-positive β-cells, following 72 hr treatment with 14-3-3i or Harmine (10μM each). (H) Cell death, as defined by the percentage of propidium iodide- (PI, 0.5μg/ml) and Hoechst 33342-(50ng/ml) positive cells, was measured in dispersed mouse islets exposed to DMSO or 14-3-3 inhibitors (10μM each) for 72 hrs. (n= 3 per group; *p < 0.05).

One of the key events underlying GSIS is the generation of ATP in mitochondria, which is needed to propagate downstream signals involved in insulin release (Hou et al., 2009). In plant cells and mouse platelets, 14-3-3 proteins have been shown to inhibit mitoplast ATP Synthase activity and mitochondrial reserve capacity, respectively (Bunney et al., 2001; Schoenwaelder et al., 2016), but whether this occurs in mouse or human β-cells is not known. Inhibition of 14-3-3 proteins in intact mouse and human islets increased glucose-induced mitochondrial activity, as measured by Oxygen Consumption Rates (OCR) (Fig. 1B,C, J,K). Moreover, the increase in OCR was associated with increased ATP synthesis rates (Fig. 1D,L), which were further confirmed by biochemical measurements of total ATP in mouse and human islets exposed to low and high glucose (Fig. 1E, M). When taken together, these findings demonstrate that 14-3-3 proteins have inhibitory effects on GSIS due in part by restricting mitochondrial function and ATP synthesis.

### Pan-inhibition of 14-3-3 proteins increases β-cell proliferation

Cell type-dependent contributions of 14-3-3 proteins on proliferation have been reported. For example, in U2OS cells genetic inhibition of 14-3-3 proteins have been shown to allow cells to prematurely enter the cell cycle and proliferate (Nguyen et al., 2004). In contrast, over-expression of some 14-3-3 isoforms in cancer cells are linked to proliferation and increased cell survival (Lim et al., 2013; Murata et al., 2012; Sambandam et al., 2015; Zhang et al., 2004). To determine if 14-3-3 proteins have roles in β-cell proliferation, two independent measurements of proliferation were used. Firstly, dispersed mouse and human islets were exposed to 14-3-3i or BV02 for 72 hrs, which led to significant increases in β-cell proliferation, as measured by the percentage of Ki67- and insulin-positive cells (Fig. 1F,N). Secondly, mouse islets were incubated with 14-3-3i and harmine for 72 hrs in the presence of EdU, and flow cytometry confirmed the ability of 14-3-3i to induce β-cell proliferation (Fig. 1G). Cell type-dependent effects on cell survival have been observed following the over-expression or depletion of various 14-3-3 proteins (Datta et al., 2000; Kuzelova et al., 2009; Masters and Fu, 2001; Xing et al., 2000), and while we previously reported that 14-3-3 protein inhibition or siRNA-mediated knockdown of 14-3-3 isoforms induced apoptosis in MIN6 insulinoma cells (Lim et al., 2013), incubation of mouse islet cells with 14-3-3i or BV02 for 72 hrs did not have detrimental effects on cell viability (Fig. 1H).

### Deletion of 14-3-3ζ in murine β-cells enhances insulin secretion

We previously characterized the metabolic phenotype of systemic 14-3-3ζ knockout (KO) mice and found that systemic deletion of 14-3-3ζ was associated with improved oral glucose tolerance due to increased circulating levels of the incretin hormone GLP-1 (Lim et al., 2016). Moreover, loss of 14-3-3ζ was associated with significantly increased β-cell area, potentially due to compensatory β-cell expansion to account for decreased insulin sensitivity (Lim et al., 2016).

To better understand how 14-3-3ζ influences β-cell function, β-cell-specific 14-3-3ζKO mice (Cre+:Flox or β14-3-3ζKO) were generated by breeding 14-3-3ζ floxed mice with *Ins1*Cre^Thor^ (Fig. 2A) (Thorens et al., 2015). Levels of *Ywhaz* mRNA were significantly reduced in β14-3-3ζKO islets (Fig. 2B). In the absence of Cre recombinase, the presence of floxed alleles for *Ywhaz* did not significantly impact body weight, glucose tolerance, or insulin secretion in male or female mice (Supp. Fig. 2A-F). No differences in intraperitoneal glucose tolerance or insulin sensitivity were detected in β14-3-3ζKO mice (Fig. 2C, D). Following an intraperitoneal glucose bolus, significantly enhanced insulin secretion was observed in β14-3-3ζKO mice, and no differences in plasma glucagon levels were detected (Fig. 2E, F). The observed enhancement in insulin secretion *in vivo* was not observed in female β14-3-3ζKO mice (Supp. Fig. 2H). No differences in β-cell mass or islet size were observed between WT or β14-3-3ζKO mice (Fig. 2G,H), but significantly increased β-cell proliferation, as measured by PCNA-positive β-cells, was observed in β14-3-3ζKO mice (Fig. 2I). Deletion of 14-3-3ζ in β-cells did not induce apoptosis (Fig. 2J).

**Figure 2.**
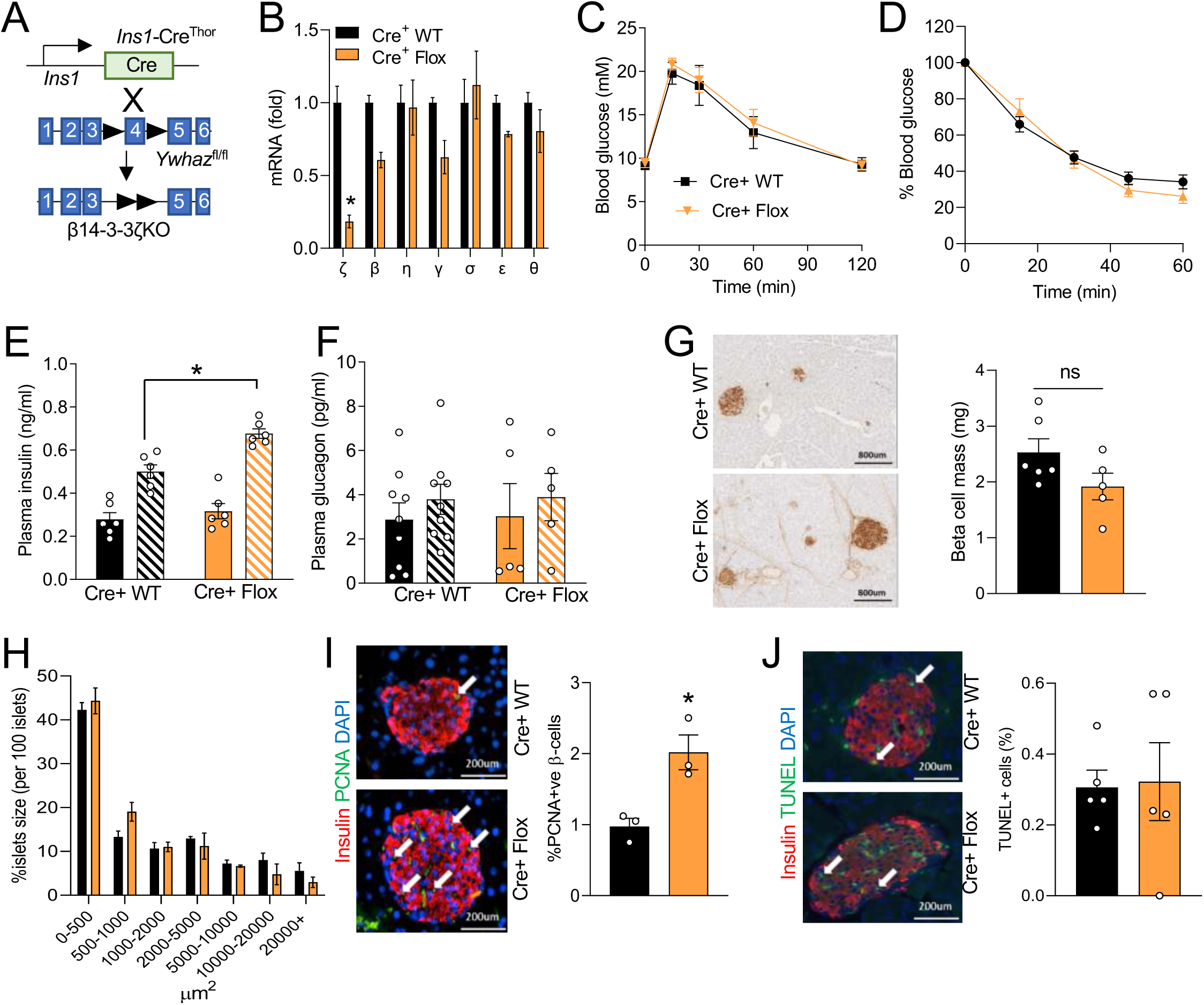
β-cell-specific deletion of 14-3-3ζ enhances glucose-induced insulin secretion *in vivo* and increases β-cell proliferation. **(A)** Generation of β-cell specific 14-3-3 KO mice (Cre^+^ Flox) was accomplished by breeding *Ins1*Cre^Thor^ mice with mice harboring floxed alleles of *Ywhaz*. **(B)** Isolated mRNA from islets from Cre^+^ Flox mice and their littermate controls (Cre^+^ WT) were subjected to qPCR analysis for 14-3-3 isoform expression. **(C,D)** No differences in glucose (C) and insulin (D) tolerance were observed in Cre+ Flox mice following intraperitoneal injections of glucose (2g/kg) or insulin (0.75 IU/kg), respectively. **(E,F)** Cre^+^ Flox mice displayed potentiated insulin secretion (E) following *i.p* glucose (2g/kg) injections and no differences were observed in circulating glucagon (F). **(G,H)** Pancreatic tissue from 12-week-old Cre^+^ WT and Cre^+^ Flox mice were collected and β-cell mass (G) and islet size distribution (H) were determined (n = 5-6 mice, four sections per mouse). **(I,J)** β-cell proliferation (I), as defined by PCNA-positive β-cells (white arrows), is increased when 14-3-3ζ is deleted. (J) TUNEL-positive apoptotic β-cells (white arrows) were measured in 4 pancreatic sections from Cre^+^ WT and Cre^+^ Flox mice (scale bar = 200μm). ( *p < 0.05 when compared to Cre^+^ WT)

Placement of 8-week old male β14-3-3ζKO mice on a high-fat diet (HFD) for 12 weeks resulted in potentiated HFD-induced weight gain (Supp Fig. 3A, B); however the increase in HFD-induced body weight gain was not associated with worsening of glucose tolerance in β14-3-3ζKO mice (Supp. Fig. 3C, D). Similar to β14-3-3ζKO mice on a control diet, an intraperitoneal glucose bolus led to enhanced insulin secretion in HFD-fed β14-3-3ζKO mice (Fig. 2E, Supp. Fig. 3E). When compared to HFD-fed control mice, β14-3-3ζKO mice fed a HFD had significantly higher β-cell mass due in part to a higher number of larger pancreatic islets (Supp. Fig. 3F-H), and this increase in β-cell mass was due to a significantly higher number of proliferating β-cells (Supp. Fig. 3I). No differences in apoptotic, TUNEL-positive β-cells were detected (Supp. Fig. 3J).

### Deletion of 14-3-3ζ in β-cells leads to profound changes in the β-cell transcriptome

As deletion of 14-3-3ζ in β-cells led to significant changes insulin secretory function and β-cell proliferation, it suggested that 14-3-3ζ affected different pathways in the β-cell, potentially due to its ability to sequester transcription factors in the cytoplasm (Brunet et al., 1999; Brunet et al., 2002; Chow and Davis, 2000). To better understand if 14-3-3ζ deletion could impact the β-cell transcriptome, single-cell RNA-seq was used to differentiate β-cells from other endocrine cells within islets. Following quality control, we first performed unbiased dimensional reduction analysis (UMAP) to identify the number of cell populations present (Supp. Table 1). Then, using known markers, we manually assigned each cluster a cell type identity. We then confirmed that deletion of 14-3-3ζ in β-cells has no impact on key marker gene expression (Fig. 3 A,B; Supp. Fig 4A) and removed any cells derived from β14-3-3ζKO mice that had incomplete knockdown of *Ywhaz*. To further verify cell identity, we then integrated previously published work from a well-defined human islet dataset (Supp. Fig. 4C). Endocrine cells were identified by the expression of *Pcsk2* and *Chga* (Supp. Fig. 4B), and clustering of endocrine cells to their specific lineages was confirmed by the expression of *Ins1, Gcg, Sst,* and *Ppy* (Fig. 3C,D). Across all identified cell types, deletion of *Ywhaz* was restricted to β-cells (Fig. 3E).

**Figure 3.**
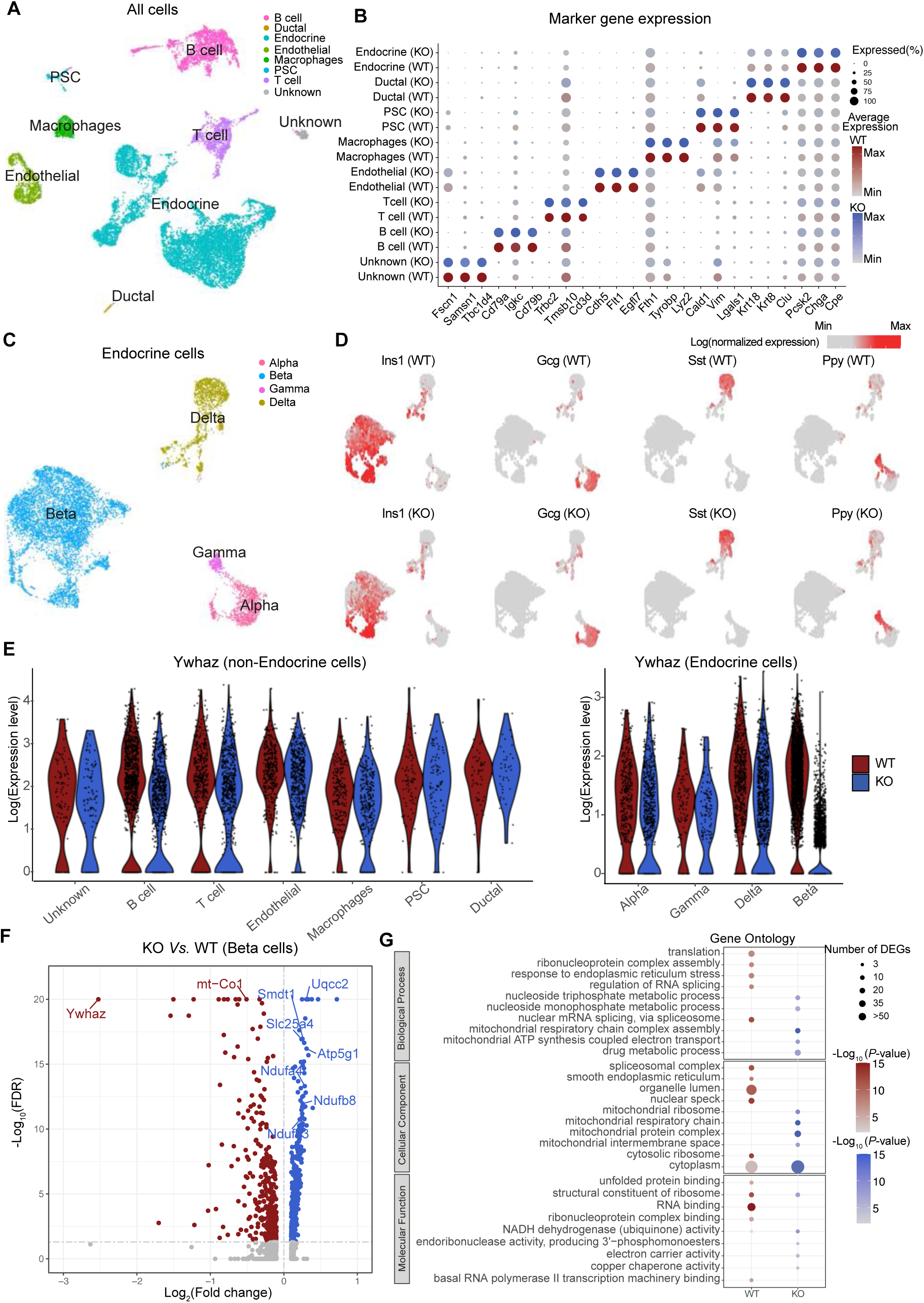
Single-cell transcriptome analyses of β14-3-3ζKO islets. **(A)** UMAP plot of all cells from wild-type and knockout mouse pancreas colored by cell type. **(B)** Dot plots of candidate marker genes specific for cell types. The size and the color of the dot encode the percentage of cells and average expression level across all cells within each group (red and blue represent high expression levels in wild-type and knockout cells, respectively). **(C)** UMAP plot of all endocrine cells from wild-type and knockout mouse pancreas colored by cell subtypes. **(D)** UMAP plot as in (C) but split into wild-type and knockout cells illustrating the expression of the four endocrine hormones: *Ins1*, *Gcg*, *Sst*, and *Ppy*. The color scale is according to log-transformed normalization values, with light grey and red colors corresponding to the minimum and maximum expression, respectively. **(E)** Violin plot showing the expression of *Ywhaz* in different types of cells. **(F)** Volcano plot of the distribution of genes with at least 0.1 difference in log2(fold change) in expression level, mapping the 351 upregulated genes (blue) and 358 downregulated genes (red). -log_10_(FDR) more than 20 was set to avoid extremely high values. **(G)** Dot plot showing enriched GO from genes highly expressed in wild-type and knockout beta cells, respectively. The size and the color of the dot encode the numbers of DEGs and the significance of corresponding enriched GO. Data are an aggregate of n=3 WT and n=3 β14-3-3ζKO mice per group.

A total of 709 differentially expressed genes (DEGs) were identified in β cells of β14-3-3ζKO mice (FDR<0.05, log_2_-fold change> 0.1, expressed in more than 15% of the cells). Of note was the discovery that among the top up-regulated DEGs in β14-3-3KO β-cells were those involved in mitochondrial metabolism, such as *Uqcc2, Atp5g1,* and *Ndufa4* (Fig. 3F,G; Supp. Table 2). Additionally, *Smdt1*, which encodes the essential regulatory subunit of the Mitochondrial Uniporter Complex (MCU), was also found to be significantly upregulated (Fig. 3F). The MCU is critical for regulating Calcium uptake into the mitochondria and for GSIS (Georgiadou et al., 2020). Gene ontology-based analysis of DEGs revealed that genes associated with regulation of RNA splicing (p= 2.7×10^-08^) were downregulated in β-cells from β14-3-3ζKO mice, which aligns with our previous finding implicating 14-3-3ζ in RNA processing and binding (Mugabo et al., 2018). Unexpectedly, enrichment of genes that participate in cell proliferation were not detected. Instead, deletion of 14-3-3ζ in β-cells led to enrichments in genes associated with mitochondrial respiratory chain complex assembly (Biological Process) and mitochondrial protein complex (Cellular Component) (p= 4.3×10^-21^ and p= 3.8×10^-28^, respectively; Fig. 3G, Supp. Fig. 4E, Supp. Table 3), which could account for the potentiation in GSIS observed *in vivo* (Fig. 2E).

### 14-3-3ζ regulates ATP-dependent insulin secretion

Deletion of 14-3-3ζ in β-cells resulted in significant changes in the expression of genes associated with mitochondrial function and respiration, key cellular processes that influence GSIS and ATP production (Fig. 3E,F). Thus, we next sought to determine whether 14-3-3ζ deletion would alter mitochondrial activity in β-cells, which could explain the enhanced GSIS in β14-3-3KO mice (Fig. 2E). Firstly, isolated islets from WT and β14-3-3ζKO mice were subjected to static GSIS assays, and under high-glucose conditions, significantly enhanced GSIS was detected from β14-3-3ζKO islets *ex vivo* (Fig. 4A). When islets from WT and β14-3-3ζKO mice were subjected to perifusion, a potentiated second-phase of insulin secretion was observed, which is consistent with the ATP-dependent phase of insulin release (Fig. 4C, D) (Henquin, 2021; Hou et al., 2009), and in the presence of diazoxide, the potentiated insulin secretory response was completely abrogated (Fig. 4C, D). Analysis of mitochondrial function and ATP synthesis revealed that 14-3-3ζ deletion in β-cells could recapitulate the augmentation in mitochondrial function and ATP synthesis following acute pan-14-3-3 protein inhibition (Fig. 4E-H). Taken together, these findings suggest that of the seven mammalian isoforms, 14-3-3ζ is likely the key mammalian isoform in β-cells that restrains insulin secretion and mitochondrial function.

**Figure 4.**
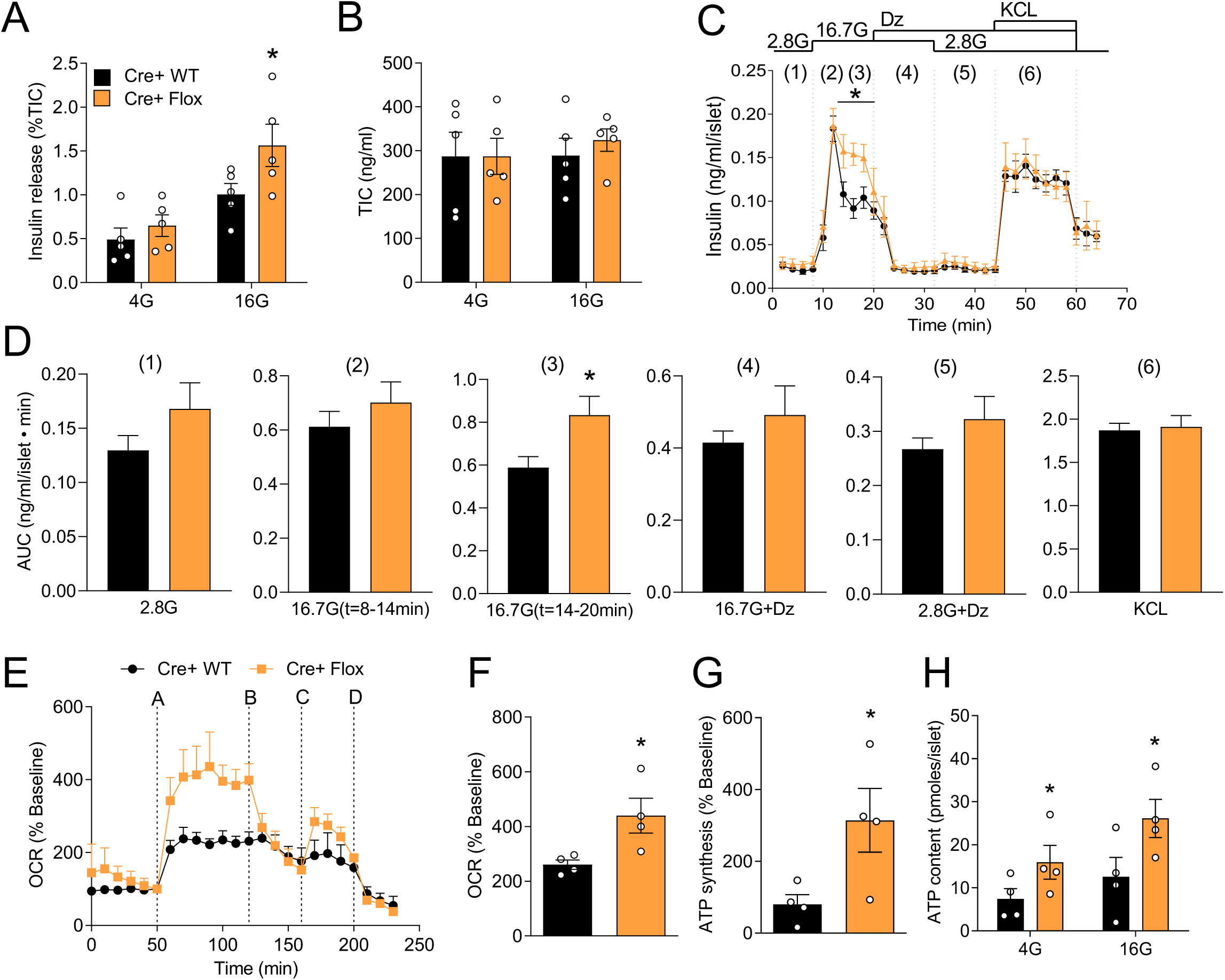
Enhanced β-cell function *ex vivo* in β-cell-specific 14-3-3ζKO mouse islets. **(A)** Isolated Islets from Cre^+^ WT and Cre^+^ Flox mice were subjected to static glucose-stimulated insulin secretion assays. **(B)** Quantification of insulin content in acid-ethanol extracts from Cre^+^ WT and Cre^+^ Flox islets (n = 5 mice per group). **(C, D)** Perifusion of islets from 14-wk-old Cre^+^ WT and Cre^+^ Flox mice was performed to examine insulin secretion dynamics in response to 16.7mM glucose, 16.7mM glucose + 200μM diazoxide, or 35mM KCl, (C) and the corresponding area under the curves (D, AUC) to each interval are shown (n = 3 mice per group). **(E-G)** Mitochondrial function, as determined by OCR, (E and F) and ATP synthesis (G) were measured in Cre^+^ WT and Cre^+^ Flox islets. **(H)** ATP content in islets from Cre^+^ WT and Cre^+^ Flox mice was quantified following exposure to different glucose concentrations (n = 4 per genotype). (*p < 0.05 when compared to Cre^+^ WT).

In addition to the upregulation of genes associated the mitochondrial respiratory chain (Fig. 3F,G) we posited that the absence of 14-3-3ζ could relieve inhibitory effects on ATP Synthase in mitochondria, which is similar to what has been observed in plants (Bunney et al., 2001). This would result in potentiated GSIS at sub-maximal glucose concentrations that would normally not induce insulin release. In this case, deletion of 14-3-3ζ would reduce the threshold needed to stimulate insulin secretion. Indeed, insulin secretion from β14-3-3ζKO mice was enhanced with 10 mM glucose (Supp. Fig. 1B). Interestingly, maximal insulin secretion in WT islets was reached at 16 mM glucose, and no further increase in insulin secretion was induced with 25 mM glucose. Although maximal GSIS from β14-3-3ζKO islets was also reached at 16 mM glucose, the magnitude of insulin release was significantly higher than in WT islets. Among the different concentrations of glucose, β14-3-3ζKO islets incubated in 10 mM glucose displayed elevations in mitochondrial activity and ATP synthesis that were equal to the maximal response of glucose (25 mM) in wildtype islets (Supp. Fig. 1B-D). Collectively, these findings demonstrate that deletion of 14-3-3ζ in β-cells lowers the glucose threshold needed for GSIS and an important role of 14-3-3ζ in regulating mitochondrial activity.

### Over-expression of 14-3-3ζ directly impairs insulin secretion and mitochondrial function

We previously reported that transgenic over-expression of TAP-14-3-3ζ in mice was associated with defective insulin secretion *in vivo*, along with impaired glucose tolerance, but it was not clear if this decrease GSIS was islet-specific (Lim et al., 2016). To further explore the negative impact of 14-3-3ζ over-expression on insulin secretion and mitochondrial function, isolated islets from WT and TAP-14-3-3ζ mice were exposed to low- and high-glucose, and in contrast to what was observed with β14-3-3ζKO islets, TAP islets demonstrated impaired GSIS (Supp. Fig. 5A). Moreover, mitochondrial function and total ATP synthesis were significantly decreased (Supp. Fig. 5D-F). To confirm whether increased 14-3-3ζ function was responsible for the impairment in GSIS, islets from TAP mice were pre-treated with 14-3-3i prior to exposure to high glucose exposure, and a restoration in GSIS comparable to WT mice was observed (Supp. Fig. 5C). Thus, over-expression of 14-3-3ζ has deleterious effects on GSIS from β-cells.

### Deletion of 14-3-3ζ influence mitochondrial dynamics in the β-cell

Using affinity proteomics, we reported interactions between 14-3-3ζ and components of ATP Synthase (Mugabo et al., 2018), and since ATP Synthase is localized to the inner mitochondrial membrane (Bunney et al., 2001; Runswick et al., 2013), it suggests that 14-3-3ζ could be mitochondrially localized. The localization of 14-3-3ζ within mitochondrial fractions has previously been reported in hippocampal neurons (Heverin et al., 2012), but it is not known if 14-3-3ζ is expressed in the mitochondrial proteome in β-cells. Purification of cytosolic and mitochondrial factors on MIN6 insulinoma cells revealed the presence of 14-3-3ζ within mitochondria (Fig. 5A), suggestive of a localized function of 14-3-3ζ.

**Figure 5.**
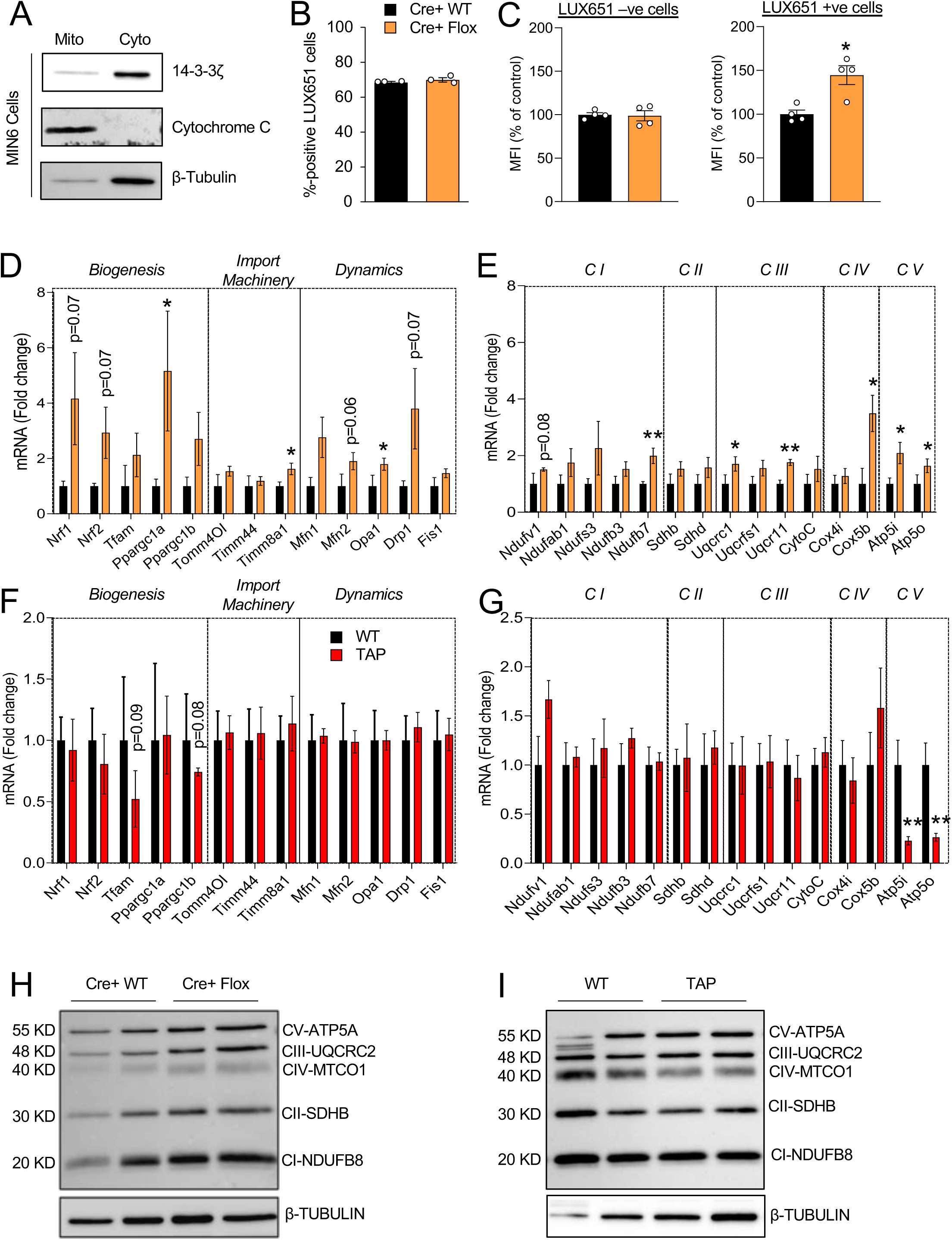
14-3-3ζ is detected in mitochondria and its deletion leads to increases in mitochondrial mass and expression of genes associated with oxidative phosphorylation and biogenesis. **(A)** Mitochondrial (Mito) and cytoplasmic (Cyto) fractions were obtained from MIN6 insulinoma cells, resolved by SDS-PAGE, and probed for 14-3-3ζ. Cytochrome C and β-tubulin were used as mitochondrial and cytoplasmic loading controls, respectively (n =3 independent experiments). **(B)** Cre^+^ WT and Cre^+^ Flox dispersed islet preparations were incubated with LUXendin651 (LUX651; 400nM) for 1 hr prior to detection by flow cytometry. The proportion of LUX651-positive cells (LUX+/total cells counted) represents β-cells from each preparation (n=4 per group). **(C)** Dispersed β14-3-3ζKO islets were treated by Mitotracker green (100nM) and LUXendin651 (400nM) to specifically label mitochondria and β-cells, respectively. Histograms depict the median fluorescent intensity (MFI) of Mitotracker green in LUX651-negative and LUX651-positive cells (n= 4 per group, *p <0.05 when compared to Cre^+^ WT). **(D-G)** Isolated mRNA from islets from Cre^+^ WT and Cre^+^ Flox mice (D,E) and WT and TAP mice (F,G) were subjected to qPCR analysis for mitochondrial biogenesis, import machinery and dynamics genes (D, F) and oxidative phosphorylation genes (E,G) (n = 3-4 mice per group, *p < 0.05; **p < 0.01 when compared to Cre^+^ WT or WT). **(H,I)** Western blot analysis of the OXPHOS mitochondrial complexes in islet extracts of Cre^+^ WT and Cre^+^ Flox mice (H; n= 3 per group) or WT and TAP mice I; (n = 3 per group).

In HCT116 cancer cells, deletion of 14-3-3σ has been found to increase mitochondrial mass (Phan et al., 2015), demonstrating the possibility that 14-3-3ζ may similarly influence in β-cell mitochondrial mass. Given the correlation between insulin secretory capacity and mitochondrial mass (Stiles and Shirihai, 2012), it raises the possibility that changes in insulin secretion following 14-3-3ζ deletion could be due to alterations in mitochondrial biogenesis. To measure mitochondrial mass in β-cells, WT and β14-3-3ζKO islets were first incubated with the fluorescent probe, LUXendin-651 (LUX651), which enriches pancreatic β-cells based upon GLP1R expression (Ast et al., 2020). In dispersed WT and β14-3-3ζKO islet preparations, no differences in the proportion of LUX651-positive cells, which represent β-cells, were detected, and approximately 70% of counted cells were LUX651-positive, which aligns with the known proportion of β-cells within a mouse islet (Fig. 5B) (Cabrera et al., 2006). Mitotracker Green was used as a surrogate measure of mass, and the median fluorescence intensity (MFI) corresponding to Mitotracker Green was significantly higher in LUX651-positive β-cells from β14-3-3ζKO islets, demonstrating increased mitochondrial mass (Fig. 5C). Notably, Mitotracker Green uptake is not dependent on mitochondrial potential. No differences in MFI were detected in LUX651-negative cells, indicating no impact on mitochondrial biogenesis in non-β-cells from β14-3-3ζKO islets (Fig. 5C). Analysis of genes related to mitochondrial biogenesis, fission, and fusion, such as *Ppgargc1a, Opa1,* and *Drp1*, were significantly increased in islets from β14-3-3ζKO mice, in addition to those regulating the mitochondrial import machinery (Fig. 5D) (Stiles and Shirihai, 2012). In contrast, no significant differences in the expression of genes associated with mitochondrial biogenesis were observed in islets from TAP mice (Fig. 5F).

As 14-3-3ζ deletion increases mitochondrial function in β14-3-3ζKO islets and is associated with enrichment of pathways associated with mitochondrial respiration within the β-cell transcriptome (Fig. 3G and 4E-G) increases in genes associated with oxidative phosphorylation, we next examined if deletion of 14-3-3ζ in β-cells could alter expression of mitochondrial complex proteins, which are necessary for ATP synthase (Vercellino and Sazanov, 2021). As measured by quantitative PCR, various genes encoding Complex proteins were significantly increased (Fig. 5E). This was in marked contrast to TAP islets, where only mRNA levels of *Atp5i* and *Atp5o* were decreased (Fig. 5G). Protein abundance of Complex proteins were also found to be elevated in β14-3-3KO mice (Fig. 5H), whereas no differences were observed in TAP islets (Supp. Fig. 6I).

### β-cell dysfunction is associated with elevated 14-3-3ζ expression and can be alleviated by 14-3-3 protein inhibition

Due to the absence of circulating leptin, *db/db* mice exhibit significant obesity develop β-cell dysfunction due to increased ER stress (Ktorza et al., 1997; Yong et al., 2021). To examine if acute 14-3-3 protein inhibition could have beneficial effects and enhance β-cell function, islets were isolated from 13 week-old male control *db/+* and diabetic, obese *db/db* mice, (Fig. 6A,B), and incubated with 14-3-3i or BV02. Similar to C57BL/6 mouse islets (Fig. 1A), enhanced GSIS was observed in both *db/+* and *db/db* mice in the presence of 14-3-3i or BV02 (Fig. 6C). Furthermore, 14-3-3 protein inhibition also resulted in enhanced mitochondrial function and ATP synthesis in *db/db* (Fig. 6D-G). To examine whether inhibition of 14-3-3 proteins could have similar effects in the context of obesity but without overt islets from 9 week-old obese *ob/ob* and non-obese *ob/+* control mice (Supp. Fig 6A,B) were isolated and exposed to 14-3-3 protein inhibitors. Similar to *db/db* mice, enhanced GSIS and mitochondrial function were observed in *ob/ob* islets following 14-3-3 protein inhibition (Supp. Fig. 6C-F).

**Figure 6.**
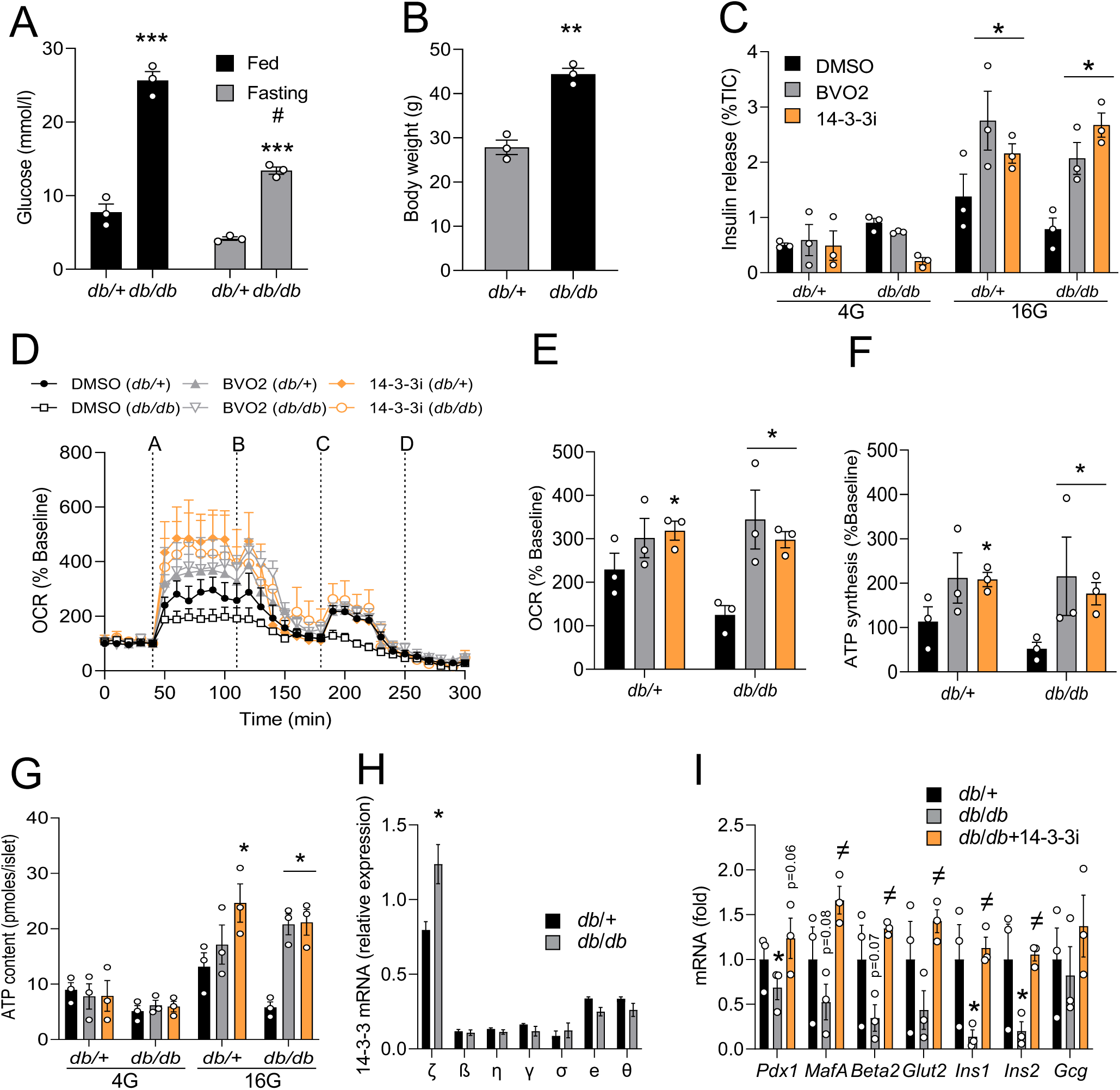
Potentiation of insulin secretion, mitochondrial function and expression of mature β-cell markers in islets from *db/db* mice. **(A,B)** Fed or fasted plasma blood glucose levels (A) and body weights **(B)** of 13-week-old *db/db* mice and control *db/+* mice (n = 3, **p <0.01; ***p <0.0001 when compared to *db/+*; # p <0.05 when compared to fasting *db/+*). **(C-F)** Islets isolated from *db/db* and *db/+* were treated with pan-14-3-3 protein inhibitors (10μM each) or DMSO, followed by static glucose-stimulated insulin secretion assays (C) or Seahorse Extracellular Flux analysis to examine mitochondrial function, as determined by oxygen consumption (OCR, D, E) and ATP synthesis (F) rates (For OCR trace: *(A)*, Glucose (16mM); *(B)*, Oligomycin (5μM); *(C)*, FCCP (1μM); *(D)*, Rotenone/Antimycin (5μM)). **(G)** ATP content of *db/db* and *db/+* mice islets treated with 14-3-3 inhibitors and quantified at different glucose concentrations. **(H and I)** Isolated mRNA from islets from *db/db* and *db/+* mice were subjected to qPCR analysis for 14-3-3 isoform expression (H) or *Pdx1, MafA, Beta2, Glut2, Ins1, Ins2 and Gcg* mRNA levels (I) (*: p < 0.05 when compared to db/+; ≠ p < 0.05 when compared to *db/db*).

Analysis of 14-3-3 isoform mRNA levels between *db/+* and *db/db* mice revealed significantly higher mRNA levels of *Ywhaz,* the gene encoding 14-3-3ζ (Fig. 6H), which suggests that elevated 14-3-3ζ expression could decrease GSIS associated with impaired mitochondrial function. In contrast, islets from *ob/ob* mice did not display differences in *Ywhaz* mRNA levels (Supp. Fig. 6G). β-cell dysfunction in *db/db* mice is associated is a loss of key genes associated with β-cell identity (Talchai et al., 2012), and when compared to *db/+* mouse islets, mRNA levels of *Ins1* and *Ins2* were significantly decreased in *db/db* islets, along with marked decreases in other mature β-cell genes like *Pdx1, Mafa,* and *Neurod1/Beta2* (Fig. 6I). Interestingly, treatment of *db/db* islets with 14-3-3i for 72 hours was sufficient to restore expression of depressed genes to levels similar to non-diabetic *db/+* islets (Fig. 6I).

Expression of *YWHAZ* mRNA in β-cells and islets from cadaveric donor islets with type 2 diabetes is increased when compared to healthy donor islets (Fig. 7A) (Segerstolpe et al., 2016). Moreover, *YWHAZ* mRNA levels were also significantly elevated in human islets from obese donors (Fig. 7A). Prolonged exposure of human islets from T2D donors to 14-3-3i or BV02 for 72 hrs stimulated the expression of genes associated with insulin biosynthesis (Fig. 7C). Similar to human islets from healthy donors, pre-treatment of islets from type 2 diabetic donors potentiated GSIS (Fig. 1I vs Fig. 7B). and 14-3-3 protein inhibition enhanced mitochondrial function and ATP synthesis in type 2 diabetic human islets (Fig. 7D-G). Taken together, these findings demonstrate the beneficial effect of inhibiting 14-3-3 proteins to enhance β-cell function in the context of overt diabetes.

**Figure 7.**
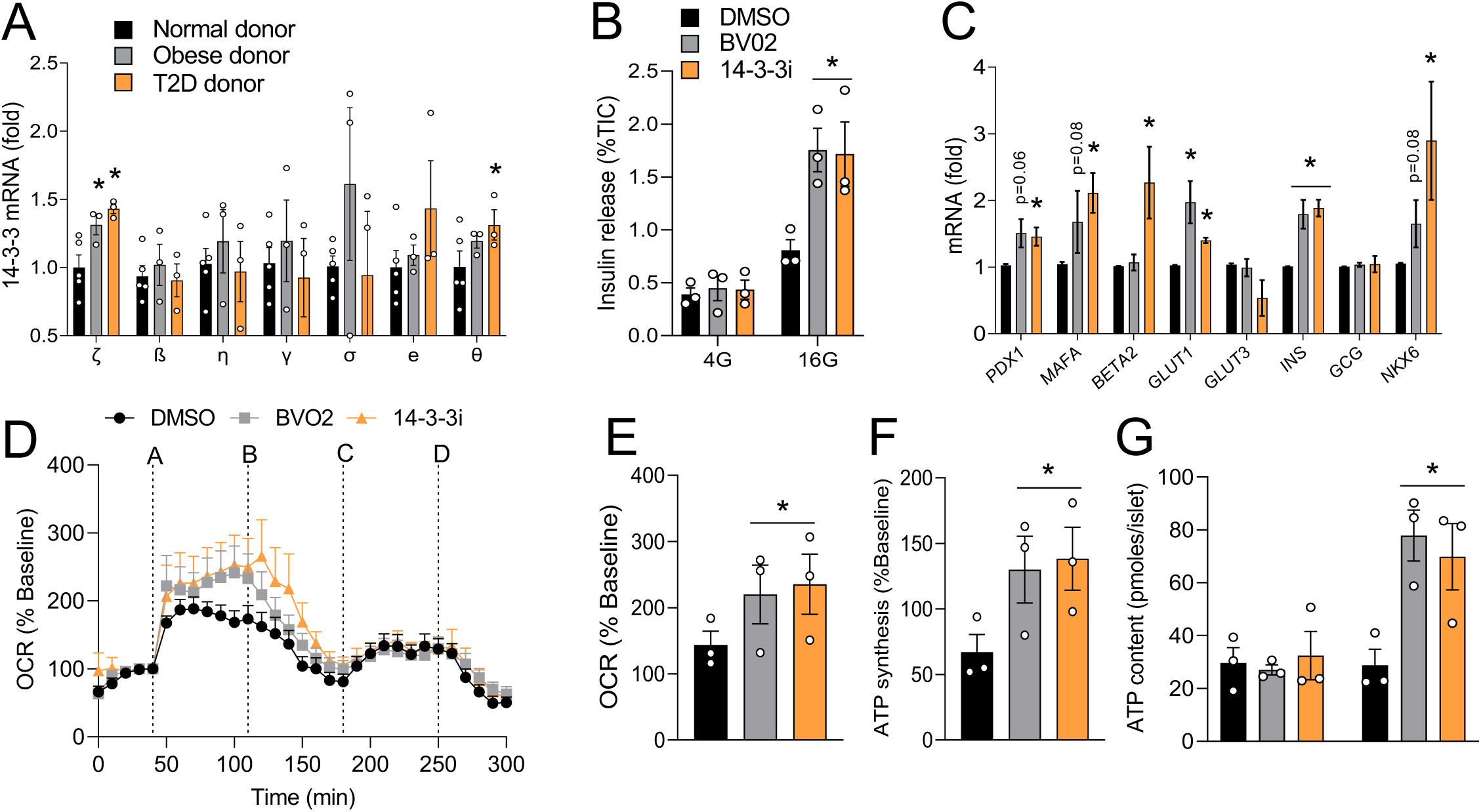
Inhibition of 14-3-3 proteins improves insulin secretory capacity and mitochondrial function in human islets from T2D donors. **(A)** Isolated mRNA from islets from normal, obese and T2D human donors were subjected to qPCR analysis for 14-3-3 isoform expression (n = 3-5 donors per group, * p <0.05 when compared to normal donor). **(B-G)** Human T2D islets (n=3 donors) were treated with pan- 14-3-3 inhibitors (10μM each) for 72hrs and a potentiation of glucose-stimulated insulin secretion was observed (B). Quantitative PCR was used to measure changes in mRNA levels of *PDX1, MAFA, BETA2, GLUT1, GLUT3, INS, GCG,* and *NKX6* after 72 hrs treatment with 14-3-3 inhibitors (10μM each). **(D-F)** T2D human islets were treated with pan-14-3-3 protein inhibitors (10μM each) or DMSO followed by Seahorse Extracellular Flux analysis to examine mitochondrial function, as determined by oxygen consumption (OCR, D, E) and ATP synthesis (F) rates (For OCR trace: *(A)*, Glucose (16mM); *(B)*, Oligomycin (5μM); *(C)*, FCCP (1μM); *(D)*, Rotenone/Antimycin (5μM)). **(G)** ATP content in T2D human islets was measured after exposure to 14-3-3 inhibitors (10μM each) and different glucose concentrations (n = 3 donors, * p < 0.05 when compared to DMSO)

## Discussion

Due to their ubiquitous expression, the importance of scaffold proteins belonging to the 14-3-3 protein family in glucose homeostasis and metabolism is often underappreciated. In the present study, we identify novel regulatory roles of 14-3-3 proteins on glucose-stimulated insulin secretion from β-cells, as they restrain insulin release due to inhibitory effects on mitochondrial function and the synthesis of ATP. Of the seven mammalian isoforms, we identify 14-3-3ζ as a key isoform that is responsible for regulating mitochondrial mass and activity and ultimately insulin secretion from the murine β-cell. Aside from effects on insulin release, inhibition of 14-3-3 proteins and deletion of 14-3-3ζ in β-cells were also found to increase β-cell proliferation. In models of β-cell dysfunction,14-3-3 protein inhibition was able to improve β-cell function and augment glucose-induced insulin release, in addition to increasing the expression of genes that regulate β-cell identity and insulin biosynthesis. These findings highlight new metabolic functions of 14-3-3ζ in β-cells and suggest the possibility of targeting 14-3-3 proteins, and specifically 14-3-3ζ, to increase β-cell function.

The signaling events underpinning glucose-stimulated insulin secretion have been well-defined, and amplifying and non-amplifying pathways have been reported (Henquin, 2000; Mugabo et al., 2017). The key organelle that is central to GSIS is the mitochondrion, which is the primary site of ATP synthesis in the β-cell. Generation of ATP requires the actions of Complex proteins (I-V) located in the inner mitochondrial membrane that work in concert to generate the proton gradient necessary for ATP synthesis (Vercellino and Sazanov, 2021). Complex V, or ATP Synthase, is the key rate-limiting enzyme involved in ATP generation (Bunney et al., 2001; Runswick et al., 2013), and inhibitory effects of 14-3-3 proteins on ATP Synthase have been reported in chloroplasts and mitoplasts (Bunney et al., 2001), and 14-3-3ζ deficiency in mouse platelets is associated with sustained intracellular ATP levels and increased mitochondrial respiratory reserve capacity (Schoenwaelder et al., 2016). To date, direct regulators of ATP synthase in β-cells have not been established. As we and others have detected 14-3-3ζ to be present within mitochondria under resting conditions (Fig. 5A) (Heverin et al., 2012) and to interact with ATP Synthase (Mugabo et al., 2018), 14-3-3ζ is likely to exert a tonic inhibitory effect on ATP synthesis within mitochondria. This may explain how acute inhibition of 14-3-3ζ and its related isoforms in mouse and human islets with 14-3-3 protein inhibitors can significantly increase mitochondrial activity and ATP synthesis. Moreover, embryonic deletion of 14-3-3ζ caused changes to the β-cell transcriptome such genes associated with oxidative phosphorylation, mitochondrial respiration, and Complex proteins were significantly upregulated, in addition to mitochondrial biogenesis. Although both acute inhibition of 14-3-3 proteins and embryonic deletion of 14-3-3ζ had different outcomes, both interventions were associated with increased mitochondrial function.

The loss of functional β-cell mass, as defined by insulin secretory capacity and β-cell number, is the key, defining feature of diabetes (Pipeleers et al., 2008; Schwartz et al., 2016). Based on results from the current study, it is tempting to speculate that targeting 14-3-3 proteins or 14-3-3ζ could have positive effects on restoring insulin secretion and β-cell mass, but more in-depth studies are needed. Expression levels of 14-3-3ζ were inversely correlated with insulin secretory capacity and mitochondrial function, as β14-3-3ζKO islets and TAP-14-3-3ζ over-expressing islets displayed potentiated or attenuated GSIS, respectively. Moreover, mRNA levels *Ywhaz/YWHAZ*, which encode for 14-3-3ζ were significantly elevated in models of β-cell dysfunction, namely *db/db* islets and human islets from type 2 diabetic donors. Due to the ubiquitous expression of 14-3-3 proteins, intensive efforts are needed to safely develop β-cell-specific approaches to target 14-3-3ζ. Interestingly, the immunomodulator FTY720 (Fingolimod), which was originally identified as a sphingosine-1-phosphate receptor agonist (Woodcock et al., 2015), has been reported to inhibit all 14-3-3 proteins by promoting the phosphorylation and subsequent dissociation of 14-3-3 protein dimers, resulting in their degradation (Woodcock et al., 2015). The effects of FTY720 on ameliorating diabetes have been documented in models of type 1 and type 2 diabetes. For example, FTY720 has been shown to delay or prevent diabetes progression in NOD mice due to effects on T cell function (Maki et al., 2002; Tsuji et al., 2012). Treatment of *db/db* mice with FTY720 reduced hyperglycemia by potently increasing β-cell regeneration when administered for more than 5 months (Zhao et al., 2012), and in spontaneously diabetic non-human primates, FTY720 administration was found to lower fasting glucose and improve β-cell function (Wang et al., 2018). Taken together, these findings suggest the future possibility of one day targeting one or more 14-3-3 isoform to treat diabetes. Alternatively, the ability to identify novel regulators of physiological pathways within the interactomes of 14-3-3ζ and its related isoforms (Dubois et al., 2009; Jin et al., 2004; Mugabo et al., 2018) presents a different 14-3-3 protein-focused approach to treat diabetes. As 14-3-3 protein inhibition had positive effects on GSIS and β-cell proliferation, it may be possible to identify novel regulators of these processes with the interactomes of 14-3-3ζ and other isoforms, and these regulators could represent new therapeutic targets for metabolic diseases.

14-3-3ζ does not have bind directly to DNA or function as a transcription factor, but deletion of 14-3-3ζ led to changes in the β-cell transcriptome. This effect is likely due to its ability to sequester transcription factors in the cytoplasm following their phosphorylation (Brunet et al., 1999; Brunet et al., 2002; Chow and Davis, 2000). Alternatively, 14-3-3 proteins may also aid in the formation of transcriptional complexes at active transcriptional “hotspots” (Siersbaek et al., 2011; Siersbaek et al., 2014). Mitochondrial DNA (mtDNA)-encoded genes are distinct from nucleus-encoded genes and are primarily transcribed and translated within mitochondria, but cooperative pathways between these two compartments are needed for proper expression of these genes (Barshad et al., 2018; Kopinski et al., 2019). This can occur through inter-organellar crosstalk between mitochondria and nuclei through shuttling of metabolites (Kopinski et al., 2019), or in the case of the present study, 14-3-3ζ could be influencing the subcellular localization of factors necessary for their expression. Prohibitin-1 and Prohibitin-2 are nuclear-encoded proteins with known roles in mitochondrial biogenesis and function, as well as the expression of Complex proteins (Ahn et al., 2006; Merkwirth et al., 2008). Moreover, deletion of Prohibitin-2 in β-cells impairs insulin secretion and mitochondrial function, causes a progressive worsening of glucose tolerance, and promotes the development of diabetes (Supale et al., 2013). In response to different stimuli, Prohibitin can translocate to different cellular compartments, including mitochondria and the nucleus (Sripathi et al., 2011). We previously identified interactions of 14-3-3ζ with Prohibitin-1 and Prohibitin-2 (Mugabo et al., 2018), and deletion of 14-3-3ζ may lead to increased abundance of Prohibitin-1 and Prohibitin-2 in mitochondria to exert positive effects on mitochondrial biogenesis or function.

β-cell proliferation is tightly regulated by the concerted nuclear actions of Cyclins and Cyclin-dependent kinases (CDKs), which are essential for the induction of proliferative genes (Cozar-Castellano et al., 2006; Hermeking and Benzinger, 2006). Moreover, various kinases, such as DYRK1A, activate parallel proliferative pathways (Dirice et al., 2016; Wang et al., 2015). Genetic inactivation of all 14-3-3 proteins has been shown to induce premature cell cycle entry and increase proliferation (Nguyen et al., 2004), and this is likely mediated by the ability of 14-3-3 proteins to regulate the activities of Cyclins and by promoting their cytoplasmic sequestration (Alvarez et al., 2007; Kim et al., 2004; Laronga et al., 2000; Lim et al., 2015) and attenuate the actions of CDK2 and CDK4 (Chan et al., 1999; Hermeking and Benzinger, 2006; Laronga et al., 2000; Nguyen et al., 2004). Alternatively, 14-3-3 proteins can influence the expression of cyclin-dependent kinase inhibitors, such as p21 and p27 (Chan et al., 1999; Hermeking and Benzinger, 2006; Laronga et al., 2000; Lim et al., 2015). Interestingly, DYRK1A is regulated by 14-3-3 proteins, as phosphorylation of DYRK1A promotes its interaction with 14-3-3 proteins and enhances its kinase activity (Alvarez et al., 2007; Kim et al., 2004). Moreover, the downstream target of DYRK1A, NFAT, is sequestered in the cytoplasm by 14-3-3 proteins when it is phosphorylated, which leads to a reduction in its transcriptional activity (Chow and Davis, 2000). Treatment of mouse and human islets with harmine and 14-3-3 inhibitors stimulated β-cell proliferation to the same extent, suggesting that the proliferative actions of 14-3-3 proteins lie upstream of DYRK1A in β-cells.

In summary, this study reports novel, physiological roles of 14-3-3 proteins in pancreatic β-cells and highlights the possibility of targeted inhibition of 14-3-3 proteins or specifically 14-3-3ζ to enhance insulin secretion. Loss of 14-3-3 protein function and deletion of 14-3-3ζ in β-cells had profound beneficial effects on mitochondrial function, glucose-stimulated insulin release, and β-cell proliferation. In contrast, increased 14-3-3ζ expression was found to be inversely associated with glucose-stimulated insulin secretion, as islets from diabetic *db/db* mice or human islets from donors with type 2 diabetes display higher levels of *Ywhaz/YWHAZ.* Moreover, inhibition of 14-3-3 proteins in these diabetic models was sufficient to enhance insulin secretion and mitochondrial function. Overall, results from the present study reveal new roles of the 14-3-3 protein family in pancreatic β-cells and deepen our understanding of the regulation of glucose-stimulated insulin release.

## Material and Methods

### Animals

All procedures were approved by the institutional committee for the protection of animals (Comité Institutionnel de Protection des Animaux du Centre Hospitalier de l’Université de Montréal). β-cell-specific 14-3-3ζ knockout mice on a C57BL/6 background and 14-3-3ζTAP transgenic mice on a CD1 background were housed on a 12-h light/dark cycle with free access to water and standard rodent chow diet (15% fat by energy). β-cell-specific 14-3-3ζ knockout mice (β14-3-3ζKO) were generated by breeding *Ins1*-Cre^Thor^ mice (JAX 026801, Thorens B. et al, 2015) with mice harboring alleles with LoxP sites flanking exon 4 of *Ywhaz*, the gene encoding 14-3-3ζ (Toronto Center for Phenogenomics, Toronto, ON) (Bradley et al., 2012; Skarnes et al., 2011). In a separate cohort, 8 week-old mice were fed with a high-fat diet (60% calories from fat, D12492, Research Diets, New Brunswick, NJ) for 12 weeks. Littermate controls (Cre+:wt/wt for β14-3-3ζ KO and wt for 14-3-3ζTAP) were used in all experiments. For glucose- and insulin-tolerance tests, β14-3-3ζKO or 14-3-3ζTAP mice were fasted for 6 h or 4h, respectively, followed by i.p. injection of 2 g/kg glucose or 0.75U/kg HumulinR insulin (Eli Lilly, Indianapolis, IN). Blood glucose levels were measured with a glucometer (Contour Next, Ascensia Diabetes Care, Mississauga, Canada). Plasma insulin and glucagon were measured by ELISA (Alpco, Salem, NH and Mercodia, NC). Male and female C57BL/KsJ *db/db* (JAX #000697) and C57BL/6J *ob/ob* (JAX #000632) mice and age-matched, lean control mice (C57BL/KsJ *db/+* or C57BL/6J *ob/+*, respectively) were purchased from the Jackson Laboratory (ME, USA).

### Mouse and human islets

Pancreatic islets were isolated by collagenase (type XI; Sigma-Aldrich, St. Louis, MO) digestion of total pancreas, as previously described (Mugabo et al., 2017). Isolated islets were handpicked under a stereoscope and cultured overnight at 37°C in 11.1 mM glucose RPMI 1640 medium with sodium bicarbonate, supplemented with 10% fetal bovine serum, 10 mM HEPES (pH 7.4), 2 mM L-glutamine, 1 mM sodium pyruvate, 100 U/ml penicillin, and 100 μg/ml streptomycin before the start of the experiments.

Human islets (75 to 90% pure; 13 different donors without any known disease and 3 type 2 diabetic donors) were from the Alberta Diabetes Institute IsletCore (Canada) and the Integrated Islet Distribution Program (City of Hope, USA). Isolated human islets were handpicked and cultured overnight in 5mM glucose DMEM medium, supplemented with 10% fetal bovine serum and 1% penicillin/streptomycin before the start of the experiments. Donor characteristics can be found in Supp. Table 4.

### Insulin secretion and ATP content measurement

Mouse islets were transferred to RPMI 1640 medium with 4 mM glucose and human islets to DMEM with 5mM glucose for 2 hours to achieve baseline insulin secretion. Batches of 10 islets for insulin secretion and 80 islets for ATP content were washed and pre-incubated for 45 minutes in Krebs Ringer buffer-Hepes (KRBH) at pH 7.4 containing 4 mM glucose and 0.5% defatted BSA and various pharmacological agents or DMSO, followed by incubation for 60 minutes in KRBH with different concentrations of glucose, in the presence or absence of pharmacological agents.

Islet perifusion (Biorep Perifusion System V5, Miami Lakes, FL) was performed to measure Insulin secretion from 80 mouse islets of equal size for each genotype. Islets were perifused at 37°C and a rate of 0.1 ml per minute with KRBH, pH 7.4, plus 0.5% BSA and glucose, diazoxide and KCl as indicated. After 20 minutes pre-perifusion in 2.8 mM glucose, the perifusate was collected every 2 minutes for 6 minutes in 2.8 mM glucose, 16 minutes in 16.7 mM glucose, 12 minutes in 16.7 mM plus 200μM diazoxide, 12 minutes in 2.8mM glucose plus 200μM diazoxide, 16 minutes in 2.8 mM glucose plus 200μM diazoxide plus 35 mM KCL and finally 6 minutes in 2.8 mM glucose. Total insulin released into medium, perifusate, and total insulin content extracted by acid ethanol were determined by radioimmunoassay (Millipore Sigma. Oakville, Canada).

Islet ATP content was determined by ATP bioluminescent assay kit (Sigma-Aldrich), as described before (Tan et al., 2019). Briefly, human and mice islets were sonicated for 1 min in 400 µL of PBS on ice. To measure the amount of ATP via luminescence, an isotonic solution containing luciferase-luciferin (LL) was prepared from a vial of ATP assay mix-lyophilized powder (Sigma-Aldrich) was used. This isotonic LL was mixed with KRBH, and ATP-dependent luciferase-luciferin (LL) bioluminescence was measured by a TD-20/20 luminometer (Turner Designs, Sunnyvale, CA). Data were recorded via Spreadsheet Interface Software v.1.2.0 (Turner BioSystems, Inc. Sunnyvale, CA).

### Oxygen consumption and mitochondrial function

Oxygen consumption was measured at 37°C from isolated mouse and human islets after overnight recovery using a Seahorse XF24 analyzer (Agilent, Santa Clara, CA). Islets were seeded at a density of 75 islets/well. After basal respiration measurement for 40 min, glucose levels were elevated to 16mM, followed by three successive injections of 5μM oligomycin, 1μM FCCP, 5μM rotenone and 5μM antimycin (Sigma-Aldrich) to assess uncoupled respiration, maximal mitochondrial respiration and non-mitochondrial respiration, respectively. ATP production was calculated by measuring the decrease in oxygen consumption rate upon injection of oligomycin (Mugabo et al., 2017).

### Quantitative Real-Time PCR

Total RNA was extracted from islet cells with the RNeasy Micro Kit (Qiagen, Hilden, Germany) and first-strand cDNA was synthesized from 0.5 to 1μg of total RNA with the High-Capacity cDNA Reverse Transcription Kit (ThermoFisher Scientific, Waltham, MA). Quantitative real-time PCR (qPCR) was performed with SYBR Green chemistry using the QuantStudio 6-flex Real-time PCR System (Thermo Fisher Scientific). Expression levels of mouse and human genes were normalized to *Actinb* or *cyclophilin* or *ACTINB*, respectively, by the ΔC_t_ method. Primer sequences are listed in Supp. Table 5.

### Immunoblotting

Cells were lysed in RIPA buffer, supplemented with protease and phosphatase inhibitors, as previously described (Lim et al., 2015; Mugabo et al., 2018). Proteins were resolved by SDS-PAGE, transferred to PVDF membranes, and probed with antibodies against 14-3-3ζ (Ab 9639), α-tubulin (Ab 2144), β-tubulin (Ab 86298), β-actin (Ab 3700) (1:1000 dilution; Cell Signaling Technology, Danvers, MA), OXPHOS complexes (110413; 1:250 dilution; Abcam, Toronto, Canada) and Cytochrome C (1:1000 dilution; BD Pharmingen, San Diego, CA, USA).

### Islet proliferation

Isolated mouse islets were dispersed in trypsin (0.05%) for 5 minutes at 37⁰C. To image proliferative β-cells, dispersed islets were seeded in chamber slides (ThermoFisher Scientific) coated with poly-D-lysine hydrobromide (Sigma-Aldrich), and cells were cultured in RPMI1640 medium containing 11 mM glucose with 10% FBS for 72 h in the presence of 10μM of BVO2, 14-3-3i and harmine (Sigma-Aldrich). Media were changed every 24 h. After successive washes with PBS, cells were fixed with 4% paraformaldehyde, followed by co-immunostaining for insulin (ab63820;1:50 dilution; Abcam) and the proliferative markers Ki67, (ab15580; 1:300; Abcam).

Human islets were handpicked, washed with PBS, and dispersed in trypsin 0.05% (ThermoFisher Scientific) for 5 min at 37°C. At the end of the digestion, cells were washed, resuspended, and plated in chamber slides pretreated with poly-D-lysine hydrobromide. After overnight incubation, dispersed human islets were incubated in DMEM with 1% fetal bovine serum for 72 h in the presence of 5 mM glucose, 10μM BVO2, 14-3-3i and harmine. The medium was changed every 24 h. At the end of treatment, cells were fixed and immunostained for insulin and Ki67. Following incubation with insulin and Ki67 antibodies, Alexa Fluor 594 and 488-conjugated secondary antibodies were used respectively (1:500 dilution; Jackson ImmunoResearch, West Grove, PA). Slides were cover-slipped after addition of ProLong Gold mounting medium (ThermoFisher Scientific). All images were taken with an Evos FL fluorescent microscope (ThermoFisher Scientific). Proliferation was calculated as the percentage of Ki67 and insulin positive cells over the total insulin positive cell population. At least 1,500 β-cells were manually counted per condition.

### Flow Cytometry

β-cell proliferation was also measured by flow cytometry. After treating with 14-3-3i or Harmine for 72 hours, islets were dispersed and dead cells were labeled using the LIVE/DEAD Fixable Aqua (405 nm) Dead Cell Stain Kit (BD Biosciences, San Jose, CA). EdU detection, using the Click-iT Plus EdU Flow Cytometry Assay Kit with Alexa Fluor 488, and immunostaining were performed according to the manufacturer’s instructions (ThermoFisher Scientific). Insulin was detected using the primary fluorophore-coupled antibody Alexa Fluor Mouse Anti-Insulin (BD Biosciences, Cat# 565689, dilution 1:50). Flow cytometry analysis was performed using a LSRIIB flow cytometer with BD FACSDiva software (BD Biosciences). Dead-cell stain, EdU-, and Insulin-labeled cells were detected using the 405-, 488-, and 640-nm lasers coupled with 525/50-, 530/30-, 670/14-nm BP filters, respectively. Proliferation was calculated as the percentage of double-positive cells for EdU and Insulin over the total Insulin-positive cell population.

To measure mitochondrial mass, 150 mouse islets from 10 week-old Cre^+^ WT or Cre^+^ Flox mice were washed PBS containing 2mM EDTA, followed by dispersion with trypsin at 37° C for 5 minutes and passed through a 30 gauge needle. Islet cells were then rinsed with PBS twice and incubated with 50nM FITC conjugated Mitotracker green (ThermoFisher Scientific) for 30 minutes at room temperature, followed by co-incubation with 400 nM LUXendin 651 (Ast et al., 2020) for an additional 30 minutes. Labelled cells were monitored for FITC (488nm/530nm) and APC (633nm⁄780nm) using flow cytometry (LSR-II, BD Biosciences) and FACSDiva software (BD Biosciences). Data were analyzed using FlowJo v10.7 (FlowJo, LLC, Ashland, USA). Median fluorescent intensity (MFI) was calculated for FITC-Mitotracker in the Luxendin 651- (FITC-Mitrotracker+/Luxendin 651-, Q1) and Luxendin 651+ (FITC-Mitrotracker +/Luxendin 651 +, Q2) populations and compared between Cre^+^ WT and Cre^+^+ Flox islets.

### Immunohistochemistry, β-cell mass measurement and TUNEL assay

Whole pancreata were removed from wild type (WT) and β14-3-3ζKO mice, weighed, fixed in 4% paraformaldehyde, embedded in paraffin and sectioned to 6-μm thickness. A minimum of 3 sections, 72 μm apart, were used in all studies. Antigen retrieval at 95°C was performed using EZ-Retriever® System (Biogenex, Fremont, CA) with 10mM sodium citrate buffer. β-cell proliferation was assessed, as described above. The In Situ Cell Death Detection Kit (TUNEL; Roche Applied Sciences, Penzberg, Germany) was used to measure β-cell apoptosis (Roche) (Lim et al., 2016). To measure β-cell mass in pancreatic sections, immunohistochemistry was performed with an insulin antibody (1:200 dilution; Ab 3014, Cell Signaling Technology, Danvers, MA) and the SignalStain® DAB Substrate Kit (Cell Signaling Technology). Hematoxylin was used for counter-staining. Slides were monitored using a high-resolution scanner (Aperio ImageScope 12.3.3) to assess the areas of insulin-positive β-cells and the whole pancreas, followed by calculating β-cell mass (total β-cell area / total pancreas weight).

### MIN6 cells culture and mitochondrial fractionation

Mouse insulinoma 6 (MIN6) cells (Dr. Jun-ichi Miyazaki, Osaka University) (passages 19–23) were cultured at 25 mM glucose in DMEM medium supplemented with 10% fetal bovine serum and 1% penicillin/streptomycin at 37°C in a humidified atmosphere (5% CO2, 95% air) (Miyazaki et al., 1990). Cells were seeded in a 75cm^2^ culture flask for 5-7 days to reach a 70–80% confluence at the day of experiment. Purification of cytoplasmic and mitochondrial fractions or pure mitochondria were done from 20 x 10^6^ cells (Mitochondria Isolation Kit for Cultured Cells, ThermoFisher).

### Single-cell RNA-seq of pancreatic islets

Islets isolated from WT and β14-3-3 KO mice at 12-14 weeks were dispersed after an overnight recovery as described above. Dead cells were removed by using a dead cell removal kit and MS columns (Miltenyi Biotec GmbH, Bergisch Gladbach, Germany). Single-cell RNA-seq libraries were made with the 10x Genomics Chromium Next GEM Single Cell 3’ Library Kit (v3.1) and sequenced using an Illumina NovaSeq6000 (100bp x2, 50k reads/cell). Approximately 9600 cells were loaded per sample with an anticipated recovery of 6000 cells.

### Reads mapping and gene expression quantification

Sequencing data were aligned and quantified by using the Cell Ranger Pipeline v6.0.2 (10x Genomics) with the default setting against the mm10 reference genome (v.2020-, downloaded from the 10X Genomics website).

### Quality control and normalization

We used 3 parameters to assess the quality of our cells. First, cut-off based on the number of expressed genes (nGene) and the expression level of mitochondrial genes (percent.mito) were used to filter out the low-quality cells. In detail, cells with percent.mito less than 0.15 and 1500 < nGene < 6000 were retained. In addition, as previously reported, reads belonging to lincRNAs Gm42418 and AY036118 were assigned toward the expression of Rn45s repeat (Kimmel et al., 2019). Cells with more than 5% of total reads mapped to Rn45s repeat are likely contaminated or dead cells (Boisset et al., 2018), as such, those cells were further removed from our analysis. Together these stringent parameters were important and included in our study considering that mitochondrial activity is impacted by 14-3-3ζ protein. Following the selection of high-quality cells, we then further filtered based on genes. Genes were considered truly expressed if they contained one or more counts in at least five cells (assessed for each sample separately). Log-normalized counts were calculated using the deconvolution strategy implemented by the computeSumFactors function in scran package (v.1.14.6) (Lun et al., 2016). We then performed rescaled normalization using the multiBatchNormfunction in the batchelor package (v.1.2.4) so that the size factors were comparable across samples (Haghverdi et al., 2018). The log-normalized expression after rescaling was used in marker gene detection and differential gene expression analysis.

### Data integration, dimensionality reduction and clustering

We integrated the filtered count matrices from the wildtype and knockout samples using the sctransform approach implemented in Seurat package (Hafemeister and Satija, 2019; Stuart et al., 2019) on the 3000 anchor features. After integration, principal component analysis (PCA) was performed on the integrated data followed by embedding into low dimensional space with Uniform Manifold Approximation and Projection (UMAP) based on the top 30 dimensions. Formed clusters generated by FindClusters function from Seurat package were assigned to cell types by consulting the expression of known marker genes and automatic annotation from SingleR package (v1.4.1) (Aran et al., 2019). To obtain the different types of endocrine cells present, cells from the endocrine cluster were further extracted and integrated based on 2000 anchor features and top 30 dimensions. In addition, scRNA-sequencing data of human pancreatic islets was integrated using the same parameters as above, with our dataset to further confirm the cell identities (Segerstolpe et al., 2016). Ortholog genes with the same gene symbols according to Ensemble annotation were used for the integration.

### Marker gene detection and differential gene expression analysis

Marker genes for each cell type were identified by FindAllMarkers function using ‘roc’ test from Seurat package. Top 50 Marker genes conserved in wild-type and mutant with at least average power more than 0.6 were selected (sup.marker.genes.tsv). Differential gene expression analysis between wild-type and knockout was performed using ‘MAST’ test implemented in FindMarkers function from Seurat package (Finak et al., 2015). Genes with an FDR less than 0.05, log_2_-(fold change) more than 0.1, and expressed in more than 15% of the cells were considered differentially expressed. The Gene Ontology (GO) enrichment analysis for differentially expressed genes was conducted using the TopGO package (Alexa, 2021). Adrian Alexi’s improved weighted scoring algorithm and Fisher’s test were used to define the significance of GO term enrichment. Functional enrichment analysis was performed using the ClusterProfile package (Yu et al., 2012). KEGG functional annotations were downloaded from EnrichR database (Kuleshov et al., 2016). Significantly enriched GO and functional terms were identified as those with a *p*-value and FDR less than 0.05, respectively. β-cells from knockout mice with expressed *Ywhaz* expression were excluded from differential gene expression analysis.

### Statistical analysis

Statistical analyses were performed through GraphPad Prism 9 by using Student’s t test or ANOVA, followed by Dunnett or Bonferroni t-tests. Statistical significance is indicated in the figures as follows: *p < 0.05; **p < 0.01; ***p < 0.001. All data are presented as mean ± SEM.

## Acknowledgements

We would like to extend our sincere thanks to donors and their families for their generosity in providing human islets, which were essential to this study. This work was supported by grants to GEL from CIHR (432626), Diabète Québec, and the Canada Research Chairs Program. Additional funding was provided by a CIHR Project Grant (156136) and a Diabetes Canada New Investigator Award to EEM. Human pancreatic islets were provided by the NIDDK-funded Integrated Islet Distribution Program (IIDP) at City of Hope, NIH Grant # 2UC4DK098085 and the JDRF-funded IIDP Islet Award Initiative. GEL holds the Canada Research Chair in Adipocyte Development. YM was previously supported by a Banting Postdoctoral Award. MG was supported by a summer studentship from Diabète Québec. SP holds the Canada Research Chair in Functional Genomics in Reproduction and Development and was also supported by the Swedish Research Council (S.P. 2016/01919), Swedish Society for Medical Research (S.P. Dnr4-236-2107) and Emil och Wera Cornells Stiftelse (S.P. Dnr4-2622/2017). DJH was supported by MRC (MR/N00275X/1 and MR/S025618/1) Project and Diabetes UK (17/0005681) Project Grants. This project has received funding from the European Research Council (ERC) under the European Union’s Horizon 2020 research and innovation programme (Starting Grant 715884 to DJH).

## Author contributions

Y.M designed and performed experiments, analyzed data, and wrote and edited the manuscript. GEL designed experiments and wrote and edited the manuscript. JJT, AG, SAC, FP, SSP, MG, and EF performed experiments. CZ analyzed data and provided bioinformatic analysis. JA and JB provided reagents. EEM, SP, and DJH edited the manuscript and provided reagents. GEL is the guarantor of this work.

## Declaration of interests

The authors declare no competing interests.

## Supplementary figures

**Supp. Figure 1.**
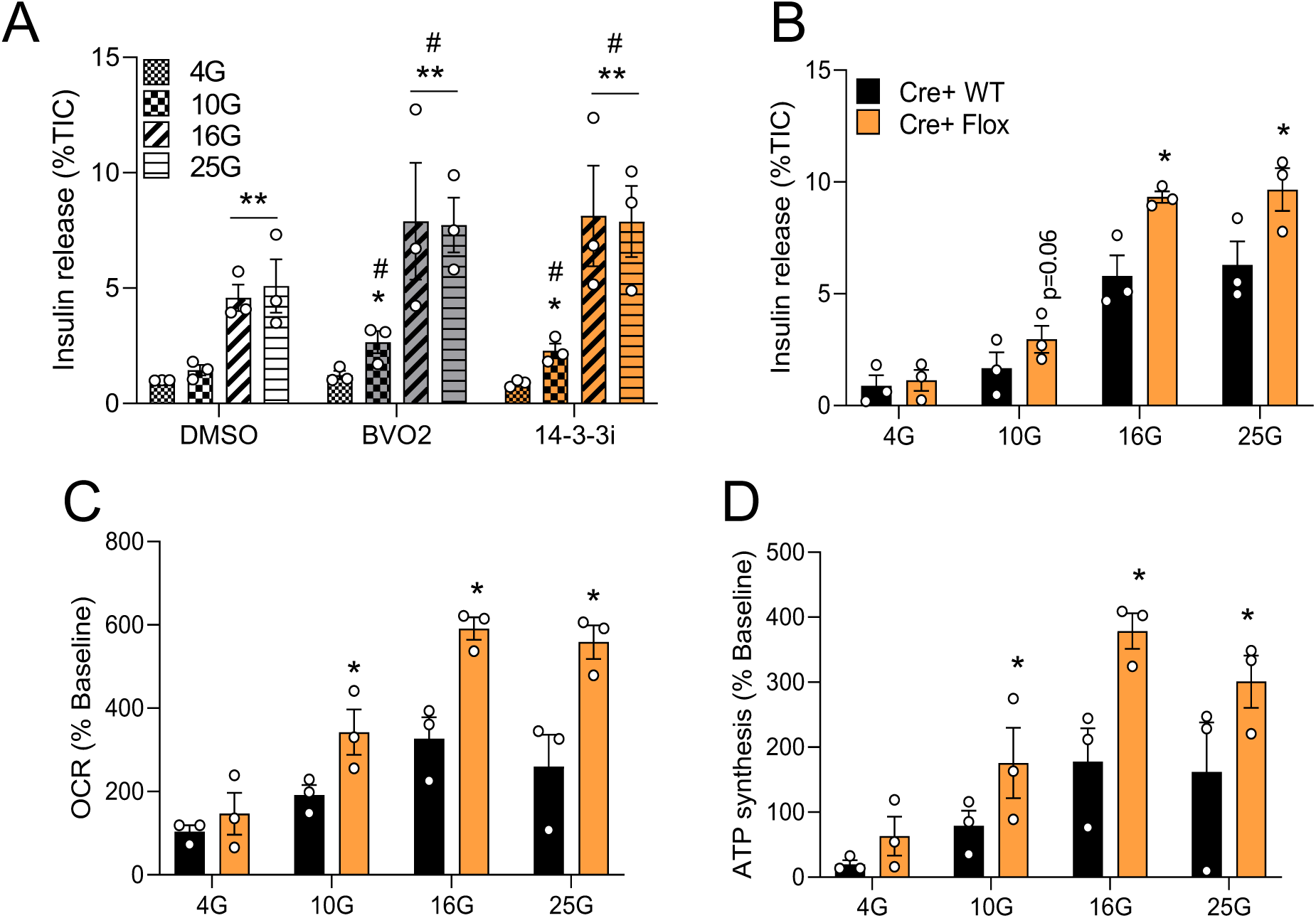
Pan-inhibition of 14-3-3 proteins or deletion of 14-3-3ζ potentiate insulin secretion and mitochondrial respiration in a glucose-dose dependent manner. **(A)** Effect of pan 14-3-3 protein inhibition on glucose-stimulated insulin secretion in mouse islets incubated at 4, 10, 16 and 25 mM glucose for 1hr (n = 3, *p < 0.05; **p <0.01 when compared to 4G; # p <0.05 when compared to DMSO). **(B-D)** Effect of β-cell-specific deletion of 14-3-3ζ on glucose-stimulated insulin secretion (B), OCR (C) and ATP synthesis (D) in islets incubated at 4, 10, 16 and 25 mM glucose for 1hr (n = 3 per group), *p <0.05 when compared to Cre^+^ WT).

**Supp. Figure 2.**
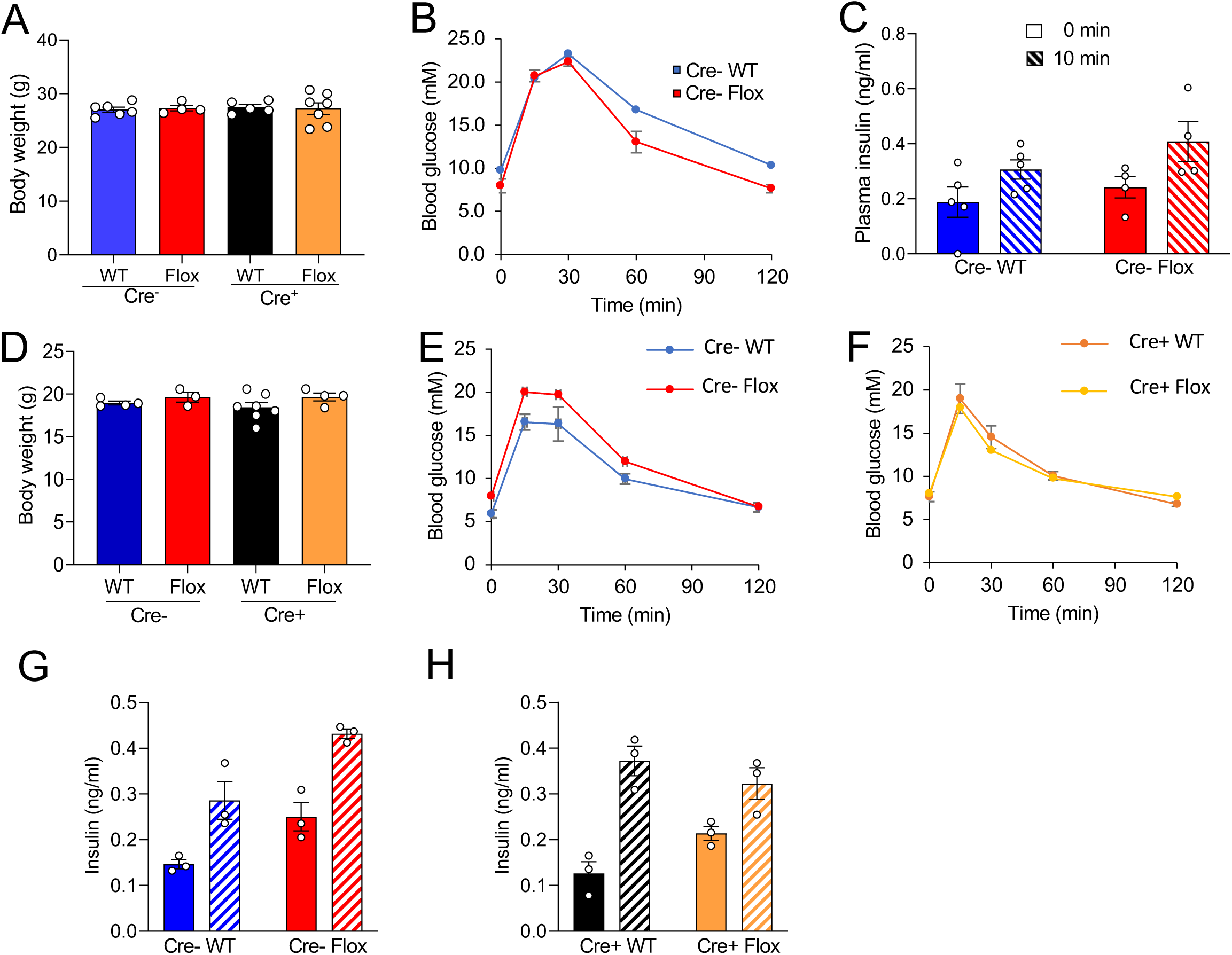
β-cell Cre expression does not affect body weight, glucose tolerance, and glucose-stimulated insulin secretion in male and female mice. **(A-C)** No differences in body weight (A) due to Cre expression were detected in male mice. Expression of Cre in β-cells did not influence glucose tolerance (B) or insulin secretion (C) following an intraperitoneal glucose (2 g/kg) bolus (n= 4-7 per group). **(D-F)** In female mice, no differences in body weight (D), glucose tolerance, (E), or insulin sensitivity (F) were detected due to the expression of Cre in β-cells. **(G,H)** No differences in glucose-stimulated insulin secretion following *i.p* glucose (2g/kg) in Cre^-^ (G) and Cre^+^ (H) female mice (n=3-7 mice per group).

**Supp. Figure 3.**
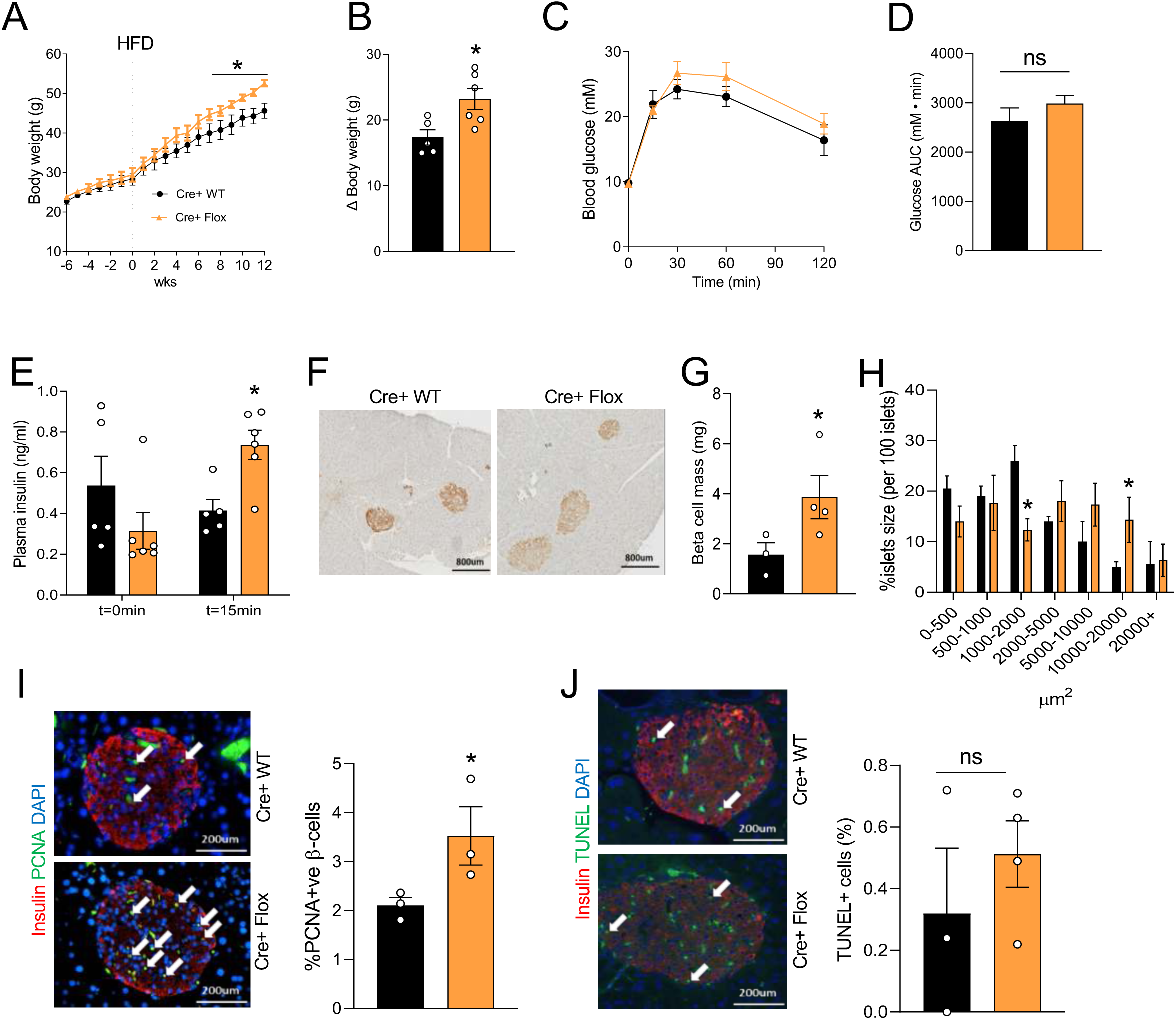
High-Fat Diet potentiates body weight gain in β-cell-specific 14-3-3ζ KO mice. **(A)** At 8 weeks of age, Cre^+^ WT and Cre^+^ Flox mice were fed a high-fat diet (HFD; 60% fat) for 12 weeks. Weekly body weights were obtained (n= 5–6 per group, *p < 0.05 when comparing to Cre^+^ WT mice). **(B)** Net body weight gain after 12 weeks of HFD feeding. **(C,D)** Intraperitoneal glucose (2g/kg) tolerance tests (C) were administered to Cre^+^ WT and Cre^+^ Flox mice after 12 weeks of HFD exposure (n= 5–6 per group). Quantification of the areas under the curve are shown in (D). **(E)** Cre^+^ Flox mice fed a HFD displayed potentiated insulin secretory response following an *i.p* glucose (2 g/kg) bolus. **(F-H)** Pancreatic tissue from HFD fed Cre^+^ WT and Cre^+^ Flox mice were collected and sectioned for the analysis. Representative islets immunohistochemistry images for the analysis of pancreatic β-cell area (F), morphometric analysis of β-cell mass (G) and islet size distribution (H) were quantified (n = 5-6 mice, four sections 100μm apart were analyzed per mouse). **(I)** β-cell proliferation, as defined by PCNA-positive β-cells (white arrows), is increased in HFD-fed Cre^+^ Flox mice when compared to Cre^+^ WT mice. Representative islet immunohistochemistry using antibodies against insulin (red), PCNA (green), and nucleus is stained blue with DAPI (n = 3 per group; four sections 100μm apart were analyzed per mouse; scale bar = 200μm; *p < 0.05 when compared to Cre^+^ WT). **(J)** β-cell apoptosis, as measured by TUNEL-positive β-cells, was determined (n = 3 per group; four sections 100μm apart were analyzed per mouse; scale bar = 200μm).

**Supp. Figure 4.**
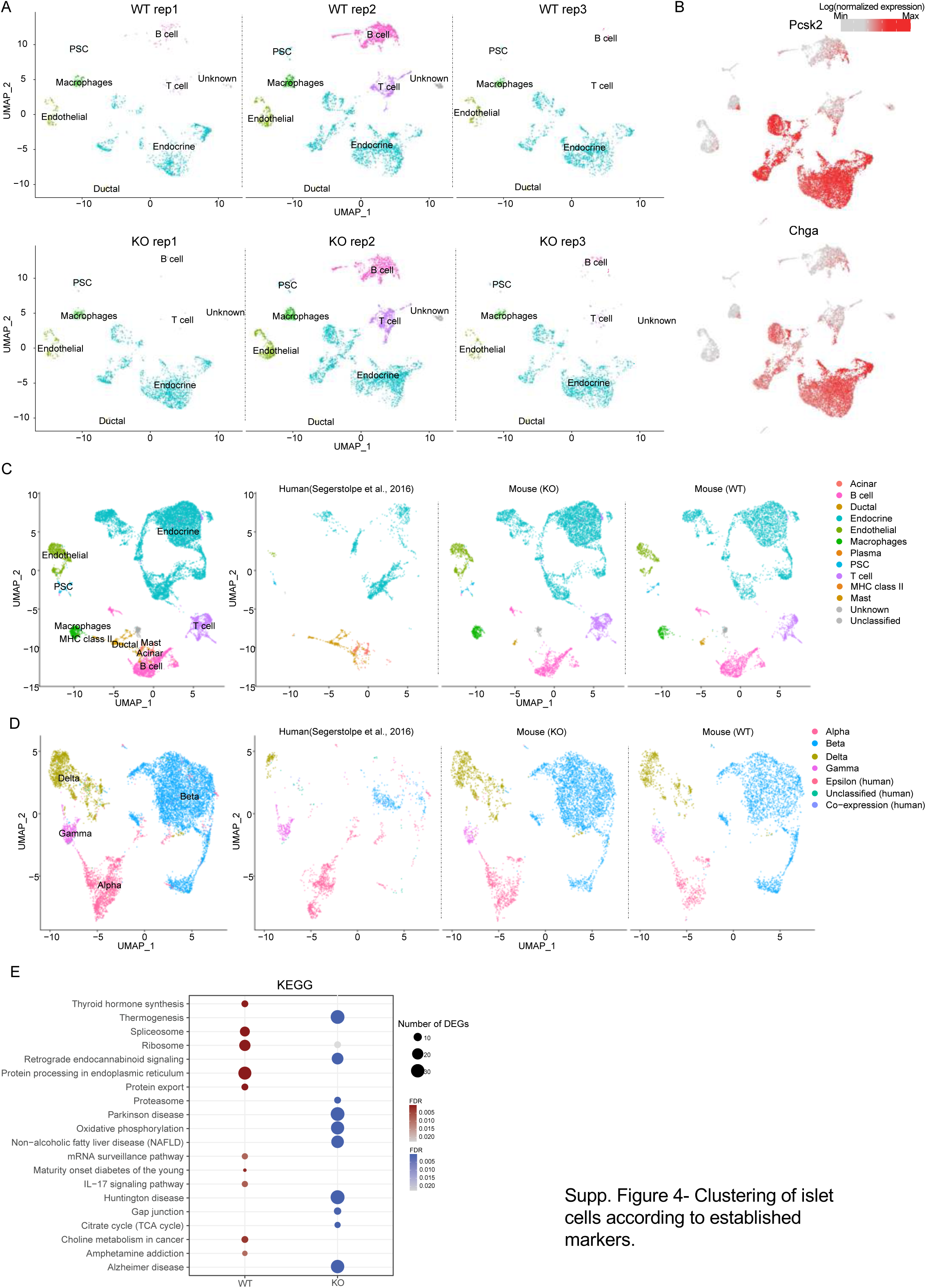
Clustering of islet cells according to established markers. **(A)** UMAP plot as in Figure 3A depicting individual samples. **(B)** Expression of endocrine marker genes (*Pcsk2* and *Chga*) overlaid onto the UMAP plot as shown in Figure 3A. **(C, D)** UMAP plot of the integrated dataset of all cells (C) and endocrine cells (D) from human pancreas dataset colored by cell types. **(E)** Dot plot showing enriched KEGG pathways from genes highly expressed in wild-type and knockout beta cells, respectively.

**Supp. Figure 5.**
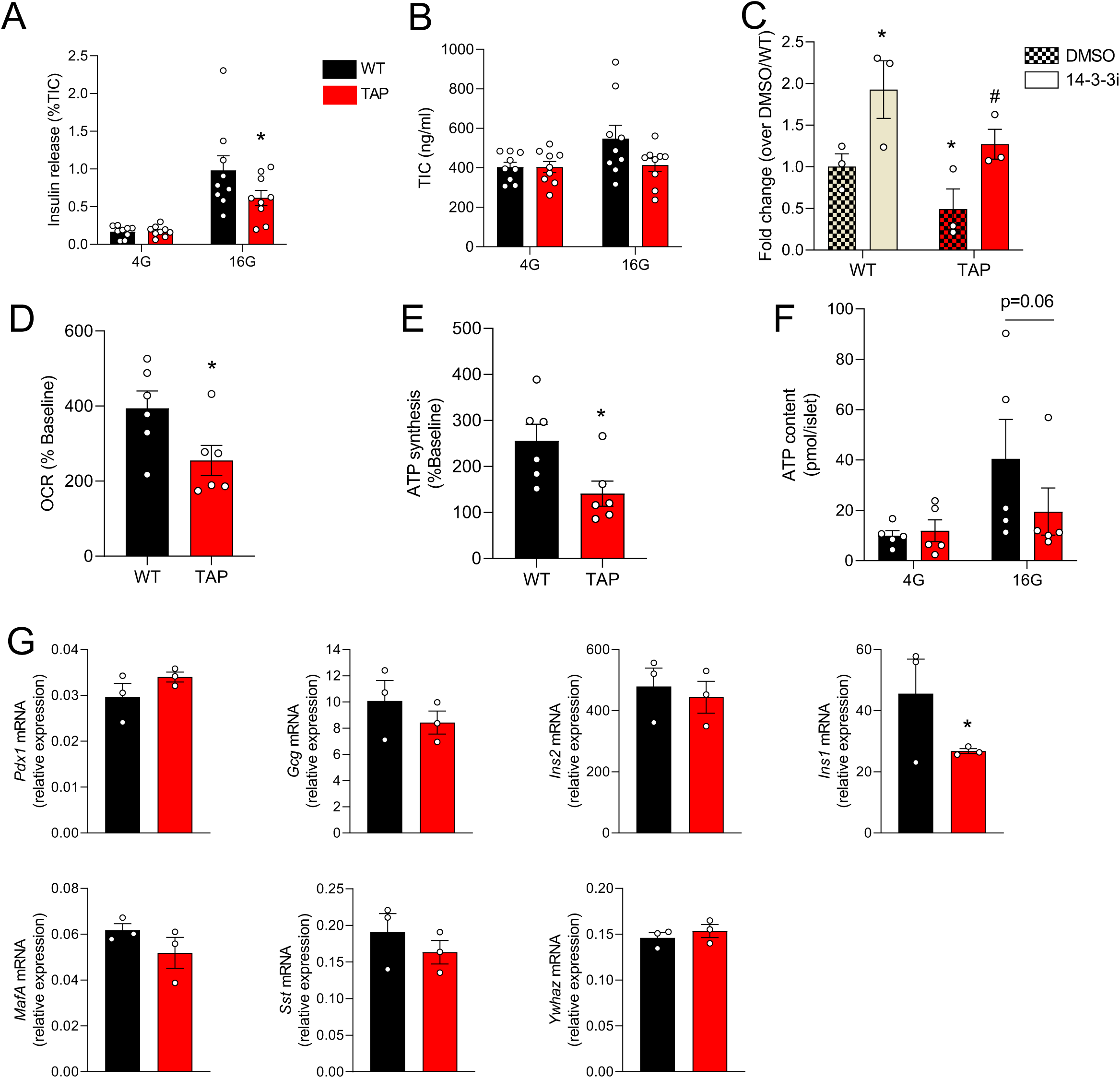
Over-expression of 14-3-3ζ in β-cells inhibits glucose-stimulated insulin secretion and mitochondrial function. **(A,B)** Islets isolated from 14-3-3ζ TAP transgenic mice and littermate WT controls were subjected to static glucose-stimulated insulin secretion assays (A). Quantification of insulin content (B) in acid-ethanol extracts from WT and 14-3-3ζTAP islets (n= 9 mice per group, *p < 0.05 when compared to WT). **(C)** Impaired GSIS due to 14-3-3ζ over-expression can be restored by pan-14-3-3 protein inhibition (*:p< 0.05 when compared to DMSO WT; # :p< 0.05 when compared to DMSO TAP). **(D, E)** Seahorse Extracellular Flux analysis to examine mitochondrial function, as determined by oxygen consumption rate (D, OCR) and ATP synthesis rates (E) (*: p< 0.05 when compared to WT). **(F)** ATP content of WT and TAP mice islets quantified at different glucose concentrations. **(G)** Isolated mRNA from islets from WT and TAP mice were subjected to qPCR analysis for *Pdx1, Gcg, Ins2, Ins1, MafA, Sst and Ywhaz* expression (n = 3 per group; *:p<0.05 when compared to WT).

**Supp. Figure 6.**
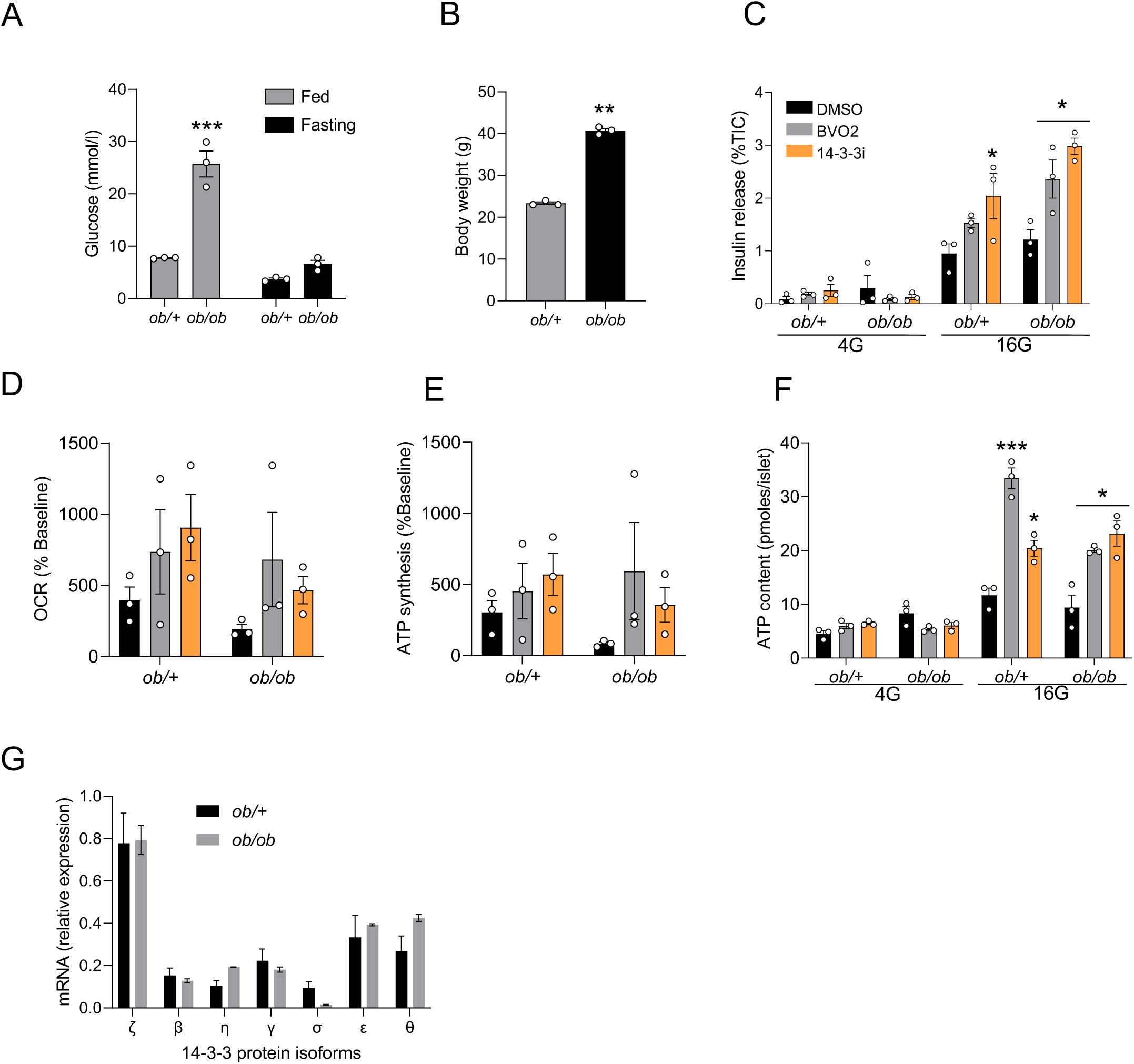
Effects of 14-3-3 protein inhibition on insulin secretion, mitochondrial function. and gene expression in islets of *ob/ob* mice. **(A, B)** Blood glucose (A) and body weights (B) of 9-week-old *ob/ob* mice and their control *ob/+* fed or fasted for 16h (n= 3; **: p <0.01; ***p <0.0001 when compared to *ob/+*). **(C-E)** Islets isolated from *ob/ob* and *ob/+* were treated with pan-14-3-3 protein inhibitors (10μM each) and subjected to static glucose-stimulated insulin secretion assays (C) and Seahorse Extracellular Flux analysis to examine mitochondrial function, as determined by OCR (D) and ATP synthesis rates (E) (*p < 0.05 when compared to DMSO). **(F)** ATP content of *ob/ob* and *ob/+* mice islets treated with 14-3-3 inhibitors and quantified at different glucose concentrations, (*p < 0.05; ***p < 0.001 when compared to DMSO). **(G)** Isolated mRNA from islets from *ob/ob* and *ob/+* mice were subjected to qPCR analysis for 14-3-3 isoform expression.

## Notes

### Competing Interest Statement

The authors have declared no competing interest.

